# Neural signature underlying the effect of intranasal vasopressin on emotional responses to spontaneous social comparison

**DOI:** 10.1101/2025.05.01.651624

**Authors:** Xiaoqing Li, Zixin Zheng, Xia Tian, Anbang Long, Binjie Yang, Ruolei Gu, Yina Ma, Wenbo Luo, Chunliang Feng

## Abstract

Vasopressin, a key molecular regulator of social behavior, is implicated in promoting self-protective responses to threats against physical safety and resources. However, its role in defending against self-view threats—common in social interactions and critical to well-being—remains unclear. This study investigates the neural mechanisms through which vasopressin modulates self-protective responses to spontaneous social comparison. In a double-blind, placebo-controlled neuroimaging experiment, participants rated their satisfaction with social evaluations and monetary outcomes assigned to a stranger, friend, or themselves. Compared to placebo, vasopressin selectively intensified contrastive emotional responses to social evaluations of the stranger, decreasing satisfaction with positive evaluations and increasing satisfaction with negative evaluations, relative to those of the self or friend. At the neural level, vasopressin reduced the distinctions between the stranger and the self/friend in the medial prefrontal cortex activity, multivariate response patterns, and functional connectivity with the temporoparietal junction and precuneus, with this effect being especially pronounced among socially dominant individuals. These converging neuroimaging findings support the hypothesis that vasopressin alters the neural representation of the stranger, shifting it from a socially irrelevant figure under placebo to a psychologically salient competitor, thereby triggering self-protective emotional processes. These findings elucidate novel neuropsychological mechanisms through which vasopressin amplifies emotional defense against social threats and offer insights into potential clinical applications for psychiatric conditions characterized by impaired self-protection.

## Introduction

Arginine vasopressin (AVP) is an evolutionarily conserved neuropeptide crucial in promoting defensive hostility against threats to survival and resources. In animals, AVP injections trigger exaggerated aggression toward strangers who pose threats to territory, mates, offspring, or physical well-being [1–5]. Conversely, blocking AVP V1a receptors (AVPR1a) attenuates both aggression and the associated autonomic responses toward intruders [3, 6]. In humans, AVPR1a genetic variations influence threat perception and avoidance [7–8], and cerebrospinal fluid AVP levels correlate with aggression history [9]. Intranasal AVP administration stimulates preemptive attacks on strangers who may threaten one’s resources [10], and elicits antagonistic responses to neutral same-sex faces of strangers as if they were threatening in men [11]. Moreover, AVP treatment and AVPR1a variations modulate brain functions in regions critical for threat detection and defensive aggression, such as the amygdala and medial prefrontal cortex (mPFC) [7, 12–15]. While the role of AVP in promoting hostility toward physical and material threats is well-established, its influence on self-protective reactions to threats to the psychological self (i.e., desired self-views) remains largely unknown. This is surprising, given that striving for desired self-views is a fundamental, uniquely human motivation that shapes various social interactions and contributes to mental health and well-being [16].

Threats to favorable self-views stem from social comparisons and evaluations that naturally occur during social interactions, often evoking defensive and hostile responses. People typically exaggerate their role in positive outcomes while deflecting blame for negative ones [17], inflate the desirability of self-views when comparing themselves/close others to an average peer or a stranger [18–19], and express aggression or denigration toward those who threaten their self-image [20–22]. These hostile self-protective behaviors are associated with contrastive emotional reactions (e.g., envy and Schadenfreude) to self-image threats in comparative interpersonal contexts [23–24]. These emotions are inherently unempathetic, marked by divergent affective responses to identical events depending on whether they affect the self or another. They reflect stronger self-protective motives [25] and mediate the impact of social comparative threats on defensive aggression [23, 26]. In short, self-image protection engages socioemotional processes closely linked to the AVP system, implicating AVP in defending against self-image threats. Accordingly, AVP elevates salivary cortisol and alters neurocognitive processes in response to social evaluative threats [27–28]. However, it remains unclear how these neurophysiological effects translate into changes in self-protective behaviors, and how AVP modulates behavioral and neural responses to social comparative threats. Our study thus investigated whether intranasal AVP amplifies contrastive emotions triggered by social comparative threats and unraveled the underlying neural mechanisms using a pharmacological fMRI approach.

The mPFC was of particular interest due to its established role in self-evaluation and self-image defense, where it encodes the personal significance of social information irrespective of valence [29]. The mPFC exhibits stronger responses to judgments about the self and close others than to strangers [30–31], shows greater activity when reflecting on the personal meaning of past experiences versus recalling concrete details [32], and tracks the importance of attributes to self-views [33]. Crucially, the assignment of personal significance by the mPFC is biased by self-protective motives, shaping behavioral responses to self-image threats. For instance, activity in the ventral mPFC (vmPFC) increases during social comparative evaluations under self-image threat, with greater activation predicting stronger defensive inflation of comparative standing [34]. The vmPFC also encodes the value of immediate self-protective decisions, while both the vmPFC and rostral mPFC (rmPFC) track defensive responses to cumulative threat [21]. Furthermore, the vmPFC and dorsal mPFC (dmPFC) are implicated in distorting threatening social evaluations to preserve desired self-views, with dmPFC activation predicting individual variability in such distortions [35–37]. Together, these findings underscore the role of mPFC subregions in representing the significance of self-image threats and orchestrating self-protective responses to perceived threats.

Building on previous work, our study examined the neuropsychological processes that mediate and modulate the AVP effect on self-protective emotional responses to social comparative threats. In a double-blind, placebo-controlled fMRI experiment, participants completed a social evaluation task where they were presented with positive or negative social evaluations received by a stranger, a close friend, or themselves, and rated their satisfaction with each outcome. They also performed a control task in which they rated satisfaction with monetary gains or losses received by each target [38]. This design allowed us to capture spontaneous responses to social comparative threats without requiring explicit social comparative judgments, and to explore whether self-protective reactions extend to close others [18]. Contrastive emotions, serving as a proxy for self-protection, were measured as differences in satisfaction ratings between the stranger and the self or friend [23]. The stronger the contrastive emotions evoked by the stranger’s outcomes, the more unsatisfied participants were with positive outcomes of the stranger compared to the self or friend, and the more satisfied they were with negative outcomes of the stranger [23–24, 26].

We hypothesized that AVP would enhance self-protective motivations by amplifying contrastive emotional responses to the stranger’s outcomes. This AVP effect was expected to be specific to social evaluations, as desired self-views are more closely tied to social feedback than to incidental monetary outcomes [39]. The AVP-induced increase in self-protective motivation would be associated with increased mPFC activity in response to evaluations of the stranger regardless of outcome valence [29], reflecting greater personal significance of the stranger under AVP treatment compared to placebo. That is, whereas the stranger may be perceived as a relatively irrelevant social target under placebo, AVP treatment may bias participants to view the stranger as a potential competitor [40], thereby increasing the self-relevance of the stranger and recruiting mPFC regions involved in self-image defense. We further hypothesized that the AVP effect would be more pronounced in socially dominant individuals. This prediction was informed by prior human and animal research demonstrating that social dominance is linked to heightened sensitivity to social comparative threats [41–42], and modulates the distribution and expression of AVP and AVPR1a to facilitate agonistic responses to social challenges [43–44]. Finally, to provide converging evidence beyond univariate activation, we explored AVP-induced alterations in multivariate neural patterns and functional connectivity of the mPFC, offering a more nuanced understanding of its influence on distributed neural processes.

In sum, this study presented a comprehensive investigation of the neuropeptidergic regulation of self-image defense, integrating behavioral and neural responses to social threat under AVP. By leveraging multiple levels of analysis, the current study aimed to provide converging evidence regarding AVP’s effects on both the psychological and neural mechanisms underlying self-image defense.

## Methods and Materials

### Participants

Forty-eight participants, comprising twenty-four pairs of male friends (*mean age* ± *SD* = 21.83 ± 3.05 years), were recruited in the study. Within each pair, one participant was randomly assigned to the intranasal AVP group (n = 24), while the other was allocated to the placebo (PBO) group (n = 24). The sample size was based on previous fMRI studies on the social functioning of AVP [28, 45] and resource constraints. The friends in each pair had known each other for more than five months and perceived each other as close friends based on their subjective ratings (see Supplementary Methods). All potential participants completed a medical history questionnaire. Potential participants were not recruited if they reported any psychiatric/clinical disorder, medication/drug/alcohol abuse, or majored in psychology/economics, or had recently participated in any other drug studies. Participants were instructed to abstain from alcohol and caffeine 2 days before the experiment and from food and drinks (except water) for 2 h before the drug administration. Two participants were excluded from further fMRI analyses due to excessive head movement during scanning (> 3 mm in translation or > 3° in rotation), resulting in 23 participants in the AVP treatment (*mean age* ± *SD* = 22.04 ± 3.80 years) and 23 participants in the PBO treatment (*mean age* ± *SD* = 21.61 ± 2.13 years). The power sensitivity analysis was conducted with MorePower software [46]. Using a 2 × 2 × 3 mixed-measures analysis of variance (ANOVA), effect sizes in the moderate-to-large range (*η^2^_p_* ≥ 0.10, repeated measures (*RM*) = 2 × 3, independent measures (*IM*) = 2, *α* = 0.05, 1 − *β* = 0.80) could be reliably detected given the current sample size [47]. The study was conducted in accordance with the 1964 Helsinki Declaration and its later amendments and was approved by the Ethics Committee of Beijing Normal University. Written informed consent was obtained from all participants before the experiment.

### Task procedure

Participants underwent two sessions on two separate days for this study (**Fig. 1**). For the first visit (see Supplementary Methods), participants were invited to the lab for a screening session about a week prior to the fMRI scanning, during which they provided sociodemographic information and completed questionnaires on medical history and personality traits. They were also asked to finish economic games, share opinions on political issues and shoot a self-introduction video. Participants were told that these materials would be sent to 8 anonymous evaluators for forming first impressions. This was to create a more reliable and engaged context to convince participants that the social evaluations they received in the subsequent social evaluation task were truly based on their personal profiles. Unbeknownst to participants, no actual evaluators were recruited, and the social evaluations assigned to different targets were predetermined [37–38]. During the second visit, participants reported their current mood states, self-administered AVP or PBO, and completed the social evaluation task and a control monetary outcome task about 35 minutes after the drug administration [11, 48]. Before leaving the lab, participants were informally debriefed, and none expressed doubts about the experimental setup. All experimental materials, including questionnaires and stimuli, were presented in the participants’ native language (i.e., Chinese).

**Fig. 1.**
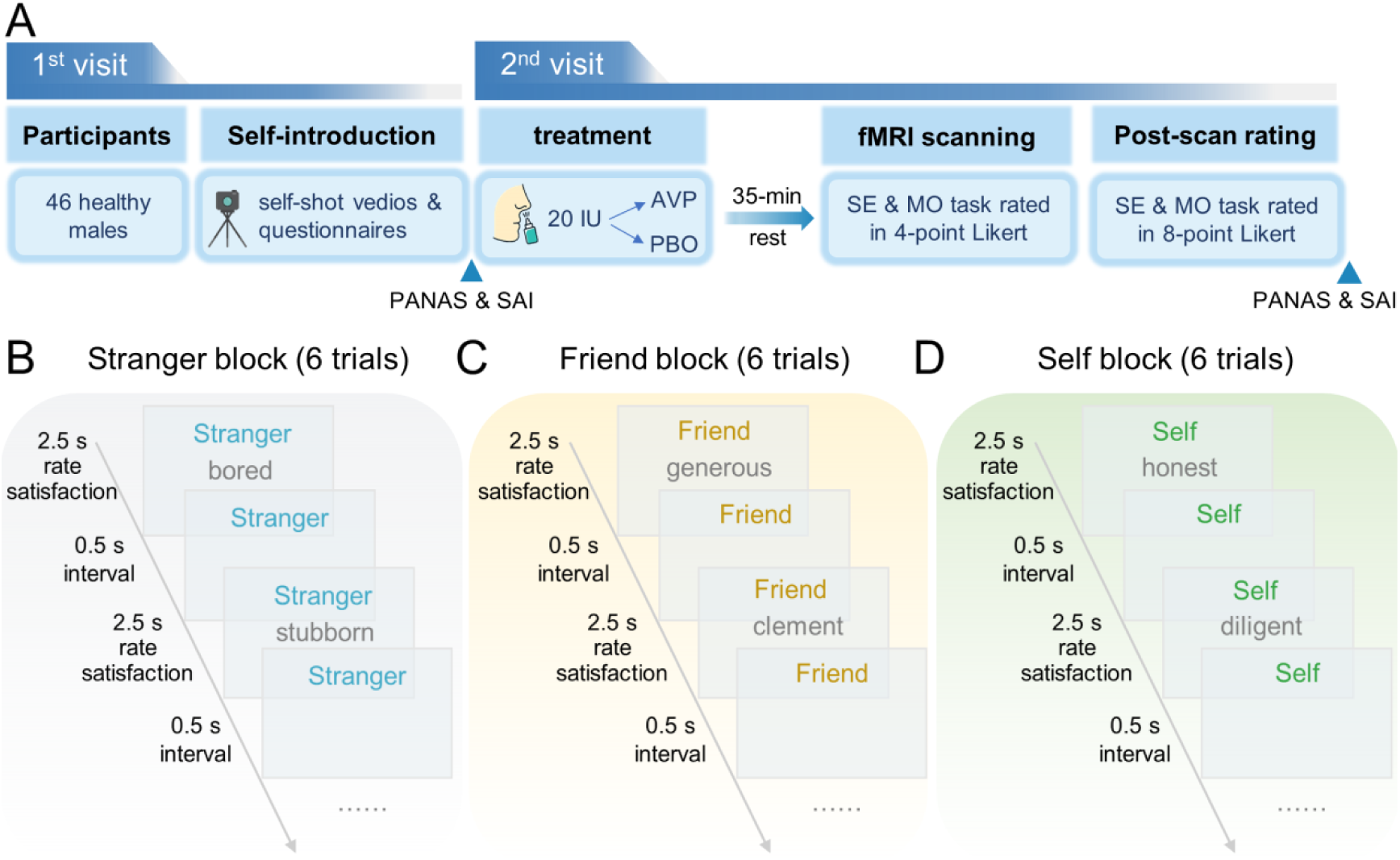
Experimental schematic. **(A)** Experimental procedure. **(B)** Timeline of the social evaluation task for a stranger block. **(C)** Timeline of the social evaluation task for a friend block. **(D)** Timeline of the social evaluation task for a self block. Each fMRI run consisted of separate blocks corresponding to six experimental conditions defined by the combination of three targets (stranger, friend, self) and two evaluation types (positive, negative). Each block began with a 3-second cue indicating the current target (“stranger,” “friend,” or “self”), followed by six consecutive evaluations directed to that target. Each evaluation was presented for 2-second followed by 0.5-second of blank screen. Two successive blocks were separated by a 10-second rest period with a central fixation. PANAS, the Positive and Negative Affect Schedule; SAI, State Anxiety Inventory; IU, international unit; AVP, arginine vasopressin; PBO, placebo; SE task, social evaluation task; MO task, monetary outcome task.

### Administration of AVP and placebo

Intranasal administration has been widely applied in humans as an effective way to cross the blood–brain barrier and directly affect central nervous system [48–49]. The AVP or PBO was randomly distributed to participants by the assistant who did not directly interact with the participants on experiment procedures, and the drug administration was blind to both participants and the experimenter. The AVP group self-administered 20 international unit (IU) of arginine vasopressin with six nasal puffs, while the PBO group self-administered six nasal puffs of sterile saline. Participants were instructed to place the nasal applicator in one nostril, and lower the lever until they felt a mist of spray in the nostril, then breathe deeply through the nose, afterwards repeat the process in another nostril. The social evaluation task and the control monetary outcome task started 35 minutes after the drug administration. After the experiment, participants were asked to report what (AVP or saline) they thought they received. The average accuracy was 50.00% for all participants, which was not significantly different from the chance level (*χ^2^*(1) = 0.00, *p* = 1.00). Moreover, there was no significant difference between PBO group and AVP group on the accuracy (*χ^2^*(1) = 0.00, *p* = 1.00).

### Social evaluation task

In the social evaluation task (**Fig. 1B**), participants were presented with positive and negative social evaluations targeting a stranger, a friend, and themselves. These evaluations were purportedly derived from eight anonymous evaluators based on personal profiles collected during the first visit. The “friend” was the individual who participated together with each participant, whereas the “stranger” was described as another anonymous participant in the study. To clarify the meaning of the “stranger” condition, participants were informed that the evaluations they and their friends received would also serve as the stranger condition for other participants. In other words, they understood that social evaluations were reciprocally exchanged among participants. Similar approaches to manipulating close others and strangers have been widely employed in research on self-relevance and self-protection [21, 35, 50].

Participants rated their satisfaction with each evaluation on a 4-point Likert scale (1 = most unsatisfied; 4 = most satisfied) during fMRI scanning and on an 8-point Likert scale (1 = most unsatisfied; 8 = most satisfied) after scanning [38]. The post-scan ratings employed a higher number of rating points compared to the during-scan ratings (8 vs. 4), a deliberate adjustment to capture participants’ satisfaction with greater precision. Rating responses were made through a response box, with associations between buttons and responses being counterbalanced across participants.

The satisfaction scale was chosen for its established utility in capturing subjective responses to both social and material outcomes [38]. Importantly, this measure aligns conceptually with the affective components of contrastive emotions, such as Schadenfreude—satisfaction derived from another’s misfortune—and envy— dissatisfaction elicited by another’s good fortune [23, 26]. Given that such socially undesirable emotions are often suppressed in explicit self-report due to impression management concerns [23, 51], participants were unlikely to report absolute satisfaction in response to negative evaluations received by the stranger and absolute dissatisfaction in response to positive ones. Instead, contrastive emotions were expected to manifest more subtly, reflected in relatively lower satisfaction with the stranger’s positive outcomes and relatively higher satisfaction with their negative outcomes, compared to those of the self or friend. Thus, it was the differential ratings between the stranger and the self/friend conditions, rather than absolute values, that were indicative of contrastive emotional responses. Consistent with the current manipulations, contrastive emotions are inherently unempathetic, characterized by markedly different affective reactions to identical events depending on whether they occur to oneself or another person [23, 51].

Importantly, the current task did not prompt explicit comparisons between targets or require direct reports of specific emotions (e.g., envy or Schadenfreude) [23]. Instead, satisfaction was rated independently for each target (self, friend, stranger) in separate blocks. This indirect approach aligns with established methods in the social comparison literature, which assess spontaneous contrastive responses through independent, target-specific judgments [18].

The social evaluation task consisted of four runs, each employing a block fMRI design. In each run, the six conditions (three targets: stranger, friend, self; two evaluation types: positive, negative) were presented in six distinct blocks, each containing six trials of the corresponding condition. The sequence of these blocks was randomized for each participant. Each block started with a 3-second cue indicating the target, followed by six evaluation trials. On each trial, participants were presented with an evaluation alongside the target cue and rated their satisfaction within 2.5 seconds followed by a 0.5-second interval. Two successive blocks were separated by a 10-second rest period with a central fixation. Stimulus presentation and behavioral data collection were implemented with Psychtoolbox-3. A total of 24 positive attributes and 24 negative attributes were employed in the social evaluation task (for a complete list of attributes, see **Table S1**).

Participants also performed four runs of monetary outcome task. In this control task (**Fig. S1**), random numbers between 10 and 99 cents were displayed as monetary gains or losses to different targets, with an average of ±54.5 cents for each condition. Participants rated their satisfaction with each lottery outcome for each target. The order of social evaluation task and monetary outcome task was counterbalanced across participants. In both the social evaluation and monetary outcome runs, three null blocks were interleaved with the six primary experimental conditions. Each of these blocks comprised six structurally identical trials associated with a given target, in which participants provided satisfaction ratings in response to a neutral label (“XXX”). These trials served to maintain attentional engagement and response consistency.

### Mood measurements

Participants filled out the Positive and Negative Affect Schedule and the State Anxiety Inventory prior to drug administration and at the end of the experiment, which respectively measured participants’ current positive affect, negative affect, and state anxiety.

### Preference for social dominance

Preference for social dominance was measured with the social dominance orientation (SDO) scale [52]. SDO scale is a widely used self-report questionnaire that measures an individual’s preference for social equality and group dominance. SDO includes 16 items and is scored on a 7-point scale from “strongly disagree” to “strongly agree”. People high in SDO typically embrace zero-sum competitive beliefs, engage more frequently in competitive social comparisons, and show stronger defensive responses to social comparative threats [53]. While the SDO scale was initially designed to measure a preference for group-level dominance, it has also been shown to capture a person’s desire for personal dominance over others [41, 54].

### fMRI data acquisition

A 3-Tesla Siemens Trio scanner was used to perform imaging, which was equipped with a twelve-channel transmit/receive gradient head coil. Functional images were acquired through a T2-weighted gradient-echo-planar imaging (EPI) sequence (repetition time [TR] = 2000 ms; echo time [TE] = 30 ms, flip angle [FA] = 90°, matrix size = 64 × 64, number of axial slices = 33, slices thickness = 4.2 mm, and field of view [FOV] = 224 mm × 224 mm). Also, a magnetization prepared rapid acquisition with gradient-echo (MPRAGE) sequence was applied to acquire high-resolution anatomical images covering the entire brain (TR = 2530 ms, TE = 3.39 ms, FA = 7°, matrix size = 256 × 256, number of slices = 144, slices thickness = 1.33 mm, FOV = 256 mm × 256 mm).

### Statistical analyses

#### Behavioral analyses

The potential differences between AVP and PBO groups in sociodemographic information, personality traits, prior- and post-experimental mood measurements were examined using two-sample t-test, Mann-Whitney U test, Chi-square test or Fisher’s precision probability test according to the types of variables, implemented in the SPSS 25.0 (IBM, Somers, USA).

The satisfaction ratings during and after fMRI scanning were analyzed with linear mixed models (LMMs), conducted with the ‘lme4’ package [55] in R version 4.2.2. Participants’ ratings were modeled with target (stranger, friend, self), type of outcomes (positive, negative), and drug treatment (AVP, PBO) as fixed variables, with participants and items included as random-effect intercept terms. The *p* values were obtained through likelihood ratio tests comparing the full model, which included the effect of interest, against a reduced model without it [56–57].

#### fMRI data analyses

Functional neuroimaging data analyses were performed with SPM12. Preprocessing of functional data included realignment through rigid-body registration to correct for head motion, spatial normalization to the Montreal Neurological Institute (MNI) template, spatial smoothing (FWHM = 8 mm) and temporal high-pass filtering.

A general linear model (GLM) was estimated for each participant with each of experimental condition as a separate regressor. The regressors were modeled using boxcar function across the corresponding blocks and convolved with the canonical hemodynamic response function (HRF). The six movement parameters of the realignment (three translations, three rotations) were also included as nuisance regressors. The resulting GLM was corrected for temporal autocorrelations using a first-order autoregressive model.

#### Univariate analyses

Regions of interest (ROIs) were created for subregions of the mPFC (dmPFC, rmPFC and vmPFC) based on a meta-analysis on personal and social reward processing across various domains [58]. These ROIs were defined as spheres with a radius of 8 mm centered at MNI coordinates of x/y/z = -10/60/26 (dmPFC), 2/56/2 (rmPFC), and 2/44/-14 (vmPFC). The coordinates were obtained by averaging all coordinates of each subregion reported in the meta-analytic study. The average parameter estimates across all voxels in each ROI were extracted from each participant for all experimental conditions using SPM Rex toolbox. These data were then analyzed using a 3 (target: stranger, friend, self) × 2 (outcome: positive, negative) × 2 (treatment: AVP, PBO) mixed-design ANOVA implemented with ‘bruceR’ package in R version 4.2.2.

The ROI analysis was complemented by an exploratory whole-brain analysis (see Supplementary Results), specifically examining the interaction between treatment and target, as consistently identified in the ROI analysis (see Results section). To control for false positives, whole-brain cluster correction was applied using a false discovery rate (FDR) threshold of *q* < 0.05 at the cluster level.

#### Moderation analyses

To test whether dispositional dominance modulates the AVP effect on the representations of social comparative threats in the mPFC, we modeled the differences in neural responses between targets (stranger vs. self, stranger vs. friend) for each mPFC subregion as a function of Treatment (AVP = 1, PBO = 0), SDO scores, and their interaction, using a linear regression model implemented with ‘bruceR’ package in R version 4.2.2.

#### Whole-brain Multi-voxel pattern analyses

*Classification for each drug treatment.* Multi-voxel pattern analysis (MVPA) was used to identify neural patterns discriminating different targets for each drug treatment, implemented with the Canlab toolbox [59]. In particular, a linear support vector machine (LSVM) was employed to train multivariate pattern classifiers to distinguish between stranger and self as well as between stranger and friend separately for AVP and PBO groups. A leave-one-out cross-validation (LOOCV) approach was employed to divide the sample into a training set (all but one participant) and a test set (the left participant) to evaluate the algorithm’s performance. The classification accuracy was evaluated using a two-choice test, where pattern expression values (i.e., the dot product of a vectorized activation image with the classifier weights) were compared for two conditions in the out-of-sample participant.

#### Comparing classification accuracy between drug treatments

The differences in classification accuracy between AVP and PBO groups were examined with permutation tests of 5,000 iterations. For each iteration, the labels of two targets were shuffled and the LOOCV approach was then employed to classify the randomly permuted targets for each drug treatment. The difference between AVP and PBO groups in the classification accuracy was computed for each iteration, resulting in the null distribution of the accuracy difference. The *p* value for the classification accuracy difference was calculated by dividing the number of models with randomly permuted targets which showed a higher accuracy difference than that of the model with true target labels by the total number of permutation (i.e., 5,000).

#### Comparing pattern expressions between drug treatments

The differences between AVP and PBO groups in multivariate neural patterns were examined by computing whole-brain pattern expressions [60]. First, the classification between targets was conducted on *N-1* participants collapsing across PBO and AVP groups, resulting in a classifier weight image for each classification. Second, pattern expression values were computed as the dot product of a vectorized activation image with the classifier weights for the two conditions tested within the same out-of-sample individual. The resultant pattern expression values were then analyzed using a mixed-design ANOVA, with target as the within-subjects factor and treatment as the between-subjects factor, implemented with ‘bruceR’ package in R version 4.2.2.

#### Cross-condition classification

The multivariate analyses revealed that AVP treatment assimilated neural responses to the stranger with those for both the self and friend (see Results section). To further support the notion that AVP produced similar effects on the stranger-self and stranger-friend distinctions, we conducted two complementary analyses. First, we collapsed the self and friend conditions and constructed a classification model to distinguish between the stranger and the combined self/friend condition for each drug treatment. Second, we performed cross-condition classification: a classifier trained to distinguish between the stranger and self was tested on the distinction between the stranger and friend in out-of-sample participants, and vice versa [61–62].

#### Determining predictive regions

To identify brain regions that robustly contributed to the classification, we employed a bootstrap test with 10,000 iterations to estimate the stability of contributing voxels. A threshold of *q(FDR)* < 0.05 at the voxel level, with a cluster size threshold (*k*) > 150, was applied to generate a thresholded weight map. In addition, we validated the contributing regions using the Haufe transformation [63]. This algorithm reconstructed activation patterns by inverting raw classifier weights to account for the covariance structure of the data. This transformation assessed the associations between each voxel and fitted values by controlling for noise and redundancy from correlated features. The same threshold was applied to the reconstructed activation patterns for the selection of contributing voxels.

#### Connectivity analyses

We examined whether mPFC worked together with other brain regions to underlie the effect of AVP on self-protection, with an analysis of psychophysiological interaction (PPI) (Friston et al., 1997), employing ROIs of dmPFC, rmPFC and vmPFC as seed regions. The fMRI signal time courses were then individually extracted from these ROIs as the seeding signals with the *generalized PPI* toolbox (McLaren et al., 2012). These seeding signals were further deconvolved with the canonical HRF, resulting in estimates of underlying neuronal activity (Gitelman et al., 2003). Subsequently, the interactions of these estimated neuronal time-series and vectors representing each of the onsets for each regressor were computed. Lastly, these interaction terms were re-convolved with the HRF and entered into a new GLM along with the vectors for the onsets of each event (i.e., the psychological terms). The contrasts between stranger and self were calculated for each participant, and the resulting contrast images were then analyzed at group level using two-sample t-tests to examine the effects of AVP treatment. For false positive control, we used whole-brain cluster correction with a cluster-level *q(FDR)* < 0.05. For the identified regions, post-hoc analyses were conducted to examine the patterns of the interaction between target (stranger, friend, self) and drug treatment (AVP, PBO).

## Results

### AVP promoted contrastive emotional responses

In the social evaluation task, the LMM analyses on the during-scan and post-scan satisfaction ratings revealed an interaction of treatment × target × outcome (during-scan ratings: *χ^2^_2_* =5.44, *p* =6.58 × 10^-2^, **Fig. 2A∼2D**; post-scan ratings: *χ^2^_2_* =34.18, *p* = 3.79 × 10^-8^, **Fig. 2E∼2H**). Follow-up analyses demonstrated that AVP decreased both during-scan and post-scan satisfaction ratings for positive evaluations received by the stranger compared to both the self (during-scan: *parameter estimates* ± *s. e.* = -0.22 ± 0.06, *p* = 7.56 × 10^-4^, **Fig. 2B**; post-scan: *parameter estimates* ± *s. e.* = -0.33 ± 0.12, *p* = 2.14 × 10^-2^, **Fig. 2F**) and friend (during-scan: *parameter estimates* ± *s. e.* = -0.21 ± 0.06, *p* = 1.13 × 10^-3^, **Fig. 2B**; post-scan: *parameter estimates* ± *s. e.* = -0.33 ± 0.12, *p* = 2.14 × 10^-2^, **Fig. 2F**).

**Fig. 2.**
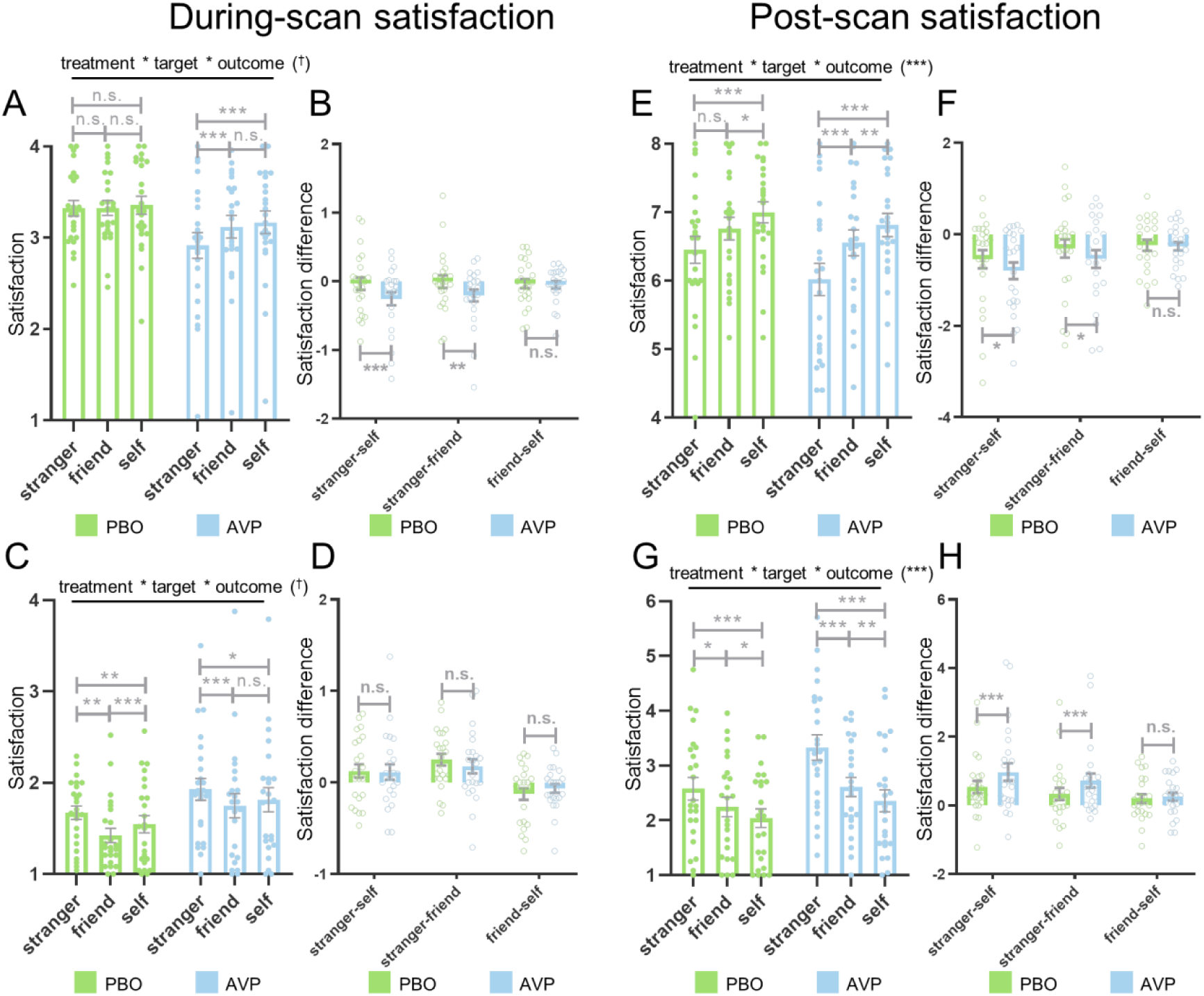
Satisfaction ratings for the social evaluation task. **(A)** During-scan satisfaction ratings for positive social evaluations as a function of treatment and target. **(B)** Pairwise comparisons of during-scan satisfaction ratings for positive social evaluations across targets as a function of treatment. **(C)** During-scan satisfaction ratings for negative social evaluations as a function of treatment and target. **(D)** Pairwise comparisons of during-scan satisfaction ratings for negative social evaluations across targets as a function of treatment. **(E)** Post-scan satisfaction ratings for positive social evaluations as a function of treatment and target. **(F)** Pairwise comparisons of post-scan satisfaction ratings for positive social evaluations across targets as a function of treatment. **(G)** Post-scan satisfaction ratings for negative social evaluations as a function of treatment and target. **(H)** Pairwise comparisons of post-scan satisfaction ratings for negative social evaluations across targets as a function of treatment. Error bars show standard error. PBO, placebo; AVP, arginine vasopressin; \*\*\**p* < 0.001; \*\**p* < 0.01; \**p* < 0.05; †*p* < 0.1; n.s., not significant.

For negative evaluations, AVP increased post-scan, but not during-scan, satisfaction ratings for outcomes assigned to the stranger compared to the self (during-scan: *parameter estimates* ± *s. e.* = -0.02 ± 0.06, *p* = 6.89 × 10^-1^, **Fig. 2D**; post-scan: *parameter estimates* ± *s. e.* = 0.55 ± 0.12, *p*= 1.41 × 10^-5^, **Fig. 2H**) and friend (during-scan: *parameter estimates* ± *s. e.* = -0.08 ± 0.06, *p* = 4.90 × 10^-1^, **Fig. 2D**; post-scan: *parameter estimates* ± *s. e.* = 0.54 ± 0.12, *p* = 1.75 × 10^-5^, **Fig. 2H**).

Notably, AVP did not significantly alter satisfaction ratings between the self and friend, either for positive (during-scan: *parameter estimates* ± *s. e.* = -0.01 ± 0.06, *p* = 8.37 × 10^-1^, **Fig. 2B**; post-scan: *parameter estimates* ± *s. e.* = -0.003 ± 0.12, *p* = 9.77 × 10^-1^, **Fig. 2F**) or negative (during-scan: *parameter estimates* ± *s. e.* = 0.06 ± 0.06, *p* = 6.39 × 10^-1^, **Fig. 2D**; post-scan: *parameter estimates* ± *s. e.* = 0.01 ± 0.12, *p* = 9.16 × 10^-1^, **Fig. 2H**) social evaluations.

In summary, AVP treatment intensified contrastive emotional responses toward social outcomes of strangers. In particular, compared to placebo, AVP reduced satisfaction ratings to the positive evaluations but enhanced satisfaction ratings to the negative evaluations received by the stranger compared to those received by the self or friend.

In the monetary outcome task, the effects pertaining to the interaction between treatment and target were not significant (see supplementary results, **Fig. S2**). The full statistical summary of satisfaction ratings is provided in **Table S2**.

### AVP enhanced mPFC responses to the social outcomes received by the stranger

In the social evaluation task, the interaction of target × treatment was significant across mPFC subregions (dmPFC: *F_2,88_* = 3.39, *p* = 3.82 × 10^-2^, *η^2^_p_* = 0.07; rmPFC: *F_2,88_* = 3.59, *p* = 3.46 × 10^-2^, *η^2^_p_* =0.08; vmPFC: *F_2,88_* =4.46, *p* = 1.69 × 10^-2^, *η^2^_p_* = 0.09; **Fig. 3A∼3C**). Under PBO treatment, activity in all mPFC subregions was lower for the stranger compared to both the self (dmPFC: *mean difference ± s. e.* = -1.80 ± 0.46, *p* = 8.54 × 10^-4^, **Fig. 3A**; rmPFC: *mean difference ± s. e.* = -2.18 ± 0.56, *p* = 9.70 × 10^-4^, **Fig. 3B**; vmPFC: *mean difference ± s. e.* = -1.40 ± 0.48, *p* = 1.56 × 10^-2^, **Fig. 3C**) and the friend (dmPFC: *mean difference ± s. e.* = -2.13 ± 0.45, *p* = 8.29 × 10^-5^, **Fig. 3A**; rmPFC: *mean difference ± s. e.* = -3.23 ± 0.70, *p*= 1.04 × 10^-4^, **Fig. 3B**; vmPFC: *mean difference ± s. e.* = -3.07 ± 0.62, *p* = 3.18 × 10^-5^, **Fig. 3C**). AVP treatment increased mPFC responses to the stranger compared to the PBO treatment (dmPFC: *mean difference ± s. e.* = 0.84 ± 0.52, *p* = 1.13 × 10^-1^, **Fig. 3A**; rmPFC: *mean difference ± s. e.* = 2.04 ± 0.59, *p* = 1.21 × 10^-3^, **Fig. 3B**; vmPFC: *mean difference ± s. e.* = 1.26 ± 0.48, *p* = 1.24 × 10^-2^, **Fig. 3C**), resulting in comparable mPFC activity between stranger and self or friend under AVP treatment (all *p* > 1.80 × 10^-1^). Other effects associated with treatment were not significant (**Table S3**).

**Fig. 3.**
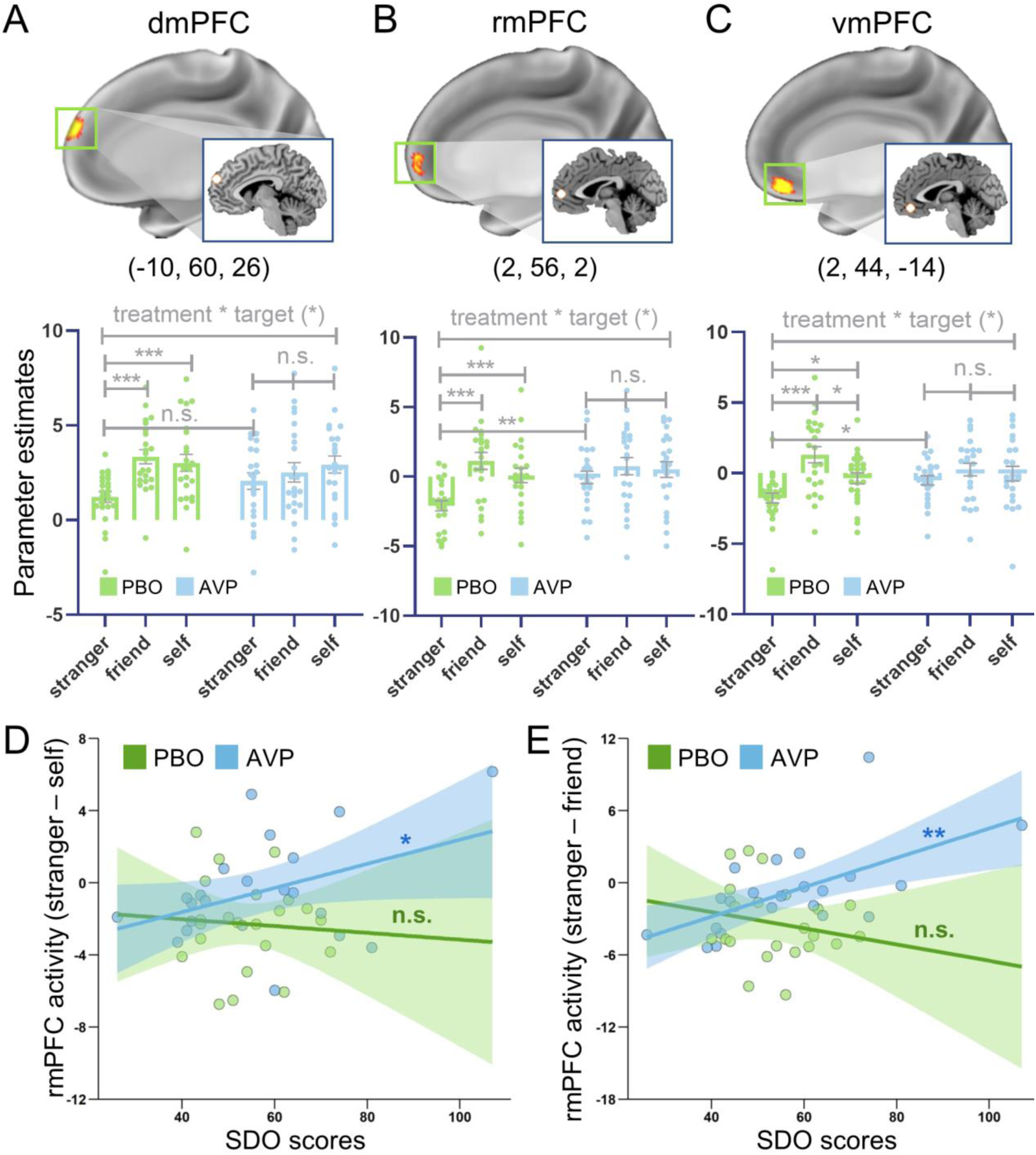
Univariate activity of mPFC subregions in the social evaluation task. **(A)** The activity of dmPFC as a function of treatment and target. **(B)** The activity of rmPFC as a function of treatment and target. **(C)** The activity of vmPFC as a function of treatment and target. **(D)** Associations between SDO scores and rmPFC activity to the contrast of stranger vs. self as a function of treatment. **(E)** Associations between SDO scores and rmPFC activity to the contrast of stranger vs. friend as a function of treatment. Blowout sections in the upper panels show sagittal slices of ROIs. Error bars show standard error. Lines represent fitted correlations, with shaded areas indicating 95% confidence intervals. dmPFC, dorsomedial prefrontal cortex; rmPFC, rostromedial prefrontal cortex; vmPFC, ventromedial prefrontal cortex; PBO, placebo; AVP, arginine vasopressin; SDO, social dominance orientation; \*\*\**p* < 0.001; \*\**p* <0.01; \**p* < 0.05; n.s., not significant.

Moreover, the drug treatment had no significant effects on mPFC responses in the monetary outcome task. A full summary of statistical results for mPFC activity is provided in **Table S3**. The results of exploratory voxel-wise whole-brain analyses are presented in the supplementary results (**Fig. S3**).

### Dispositional dominance was associated with enhanced AVP effects on rmPFC responses

The impact of AVP on rmPFC responses was stronger among individuals with higher SDO scores for the contrast between stranger and friend (*F_1, 40_* = 5.43, *p* = 2.50 × 10^-2^, **Fig. 3E**), with a similar trend for the contrast between stranger and self (*F_1, 40_* = 1.49, *p* = 2.30 × 10^-1^, **Fig. 3D**). Simple slope analyses further revealed that under AVP treatment, higher SDO scores were positively associated with increased rmPFC responses to the stranger relative to both self and friend (stranger vs. self: *β* ± *s. e.* = 0.07 ± 0.03, *p* = 4.65 × 10^-2^, **Fig. 3D**; stranger vs. friend: *β* ± *s. e.* = 0.12 ± 0.04, *p* =2.39 × 10^-3^, **Fig. 3E**). In contrast, under PBO treatment, SDO scores were not significantly associated with rmPFC activity for either contrast (stranger vs. self: *β* ± *s. e.* = -0.02 ± 0.06, *p* =7.64 × 10^-1^, **Fig. 3D**; stranger vs. friend: *β* ± *s. e.* = -0.07 ± 0.07, *p* = 3.56 × 10^-1^, **Fig. 3E**). These results were corroborated by complementary analyses collapsing self and friend conditions (see supplementary results, **Fig. S6A**). Similar patterns were observed for the dmPFC and vmPFC, though the interaction between SDO scores and drug treatment did not reach significance in these regions (see supplementary results, **Fig. S4**). Collectively, these findings suggest that individuals higher in social dominance are more susceptible to the effects of AVP.

### AVP blunted the neural discrimination between stranger and self or friend

Under the PBO treatment, multivariate neural patterns significantly distinguished the stranger from the self (*accuracy* = 100.00%, 95% *confidence interval [CI]*: 100.00 ∼ 100.00%, *discriminability [d]* = 2.00, *p* = 2.38 × 10^-7^, **Fig. 4A**) and the friend (*accuracy* = 95.65%, 95% *CI*: 87.32 ∼ 100.00%, *d* = 1.91, *p* = 5.72 × 10^-6^, **Fig. 4E**). In contrast, AVP treatment reduced these distinctions to chance levels for both the stranger-self (*accuracy* = 69.57%, 95% *CI*: 50.76 ∼ 88.37%, *d* = 0.91, *p* = 9.31 × 10^-2^, **Fig. 4B**) and stranger-friend classifiers (*accuracy* = 69.57%, 95% *CI*: 50.76 ∼ 88.37%, *d* = 0.48, *p* = 9.31 × 10^-2^, **Fig. 4F**). Moreover, classification performance was higher under the PBO compared to AVP treatment for both the stranger-self (*p* = 1.86 × 10^-2^, **Fig. 4C**) and stranger-friend distinctions (*p* = 4.98 × 10^-2^, **Fig. 4G**).

**Fig. 4.**
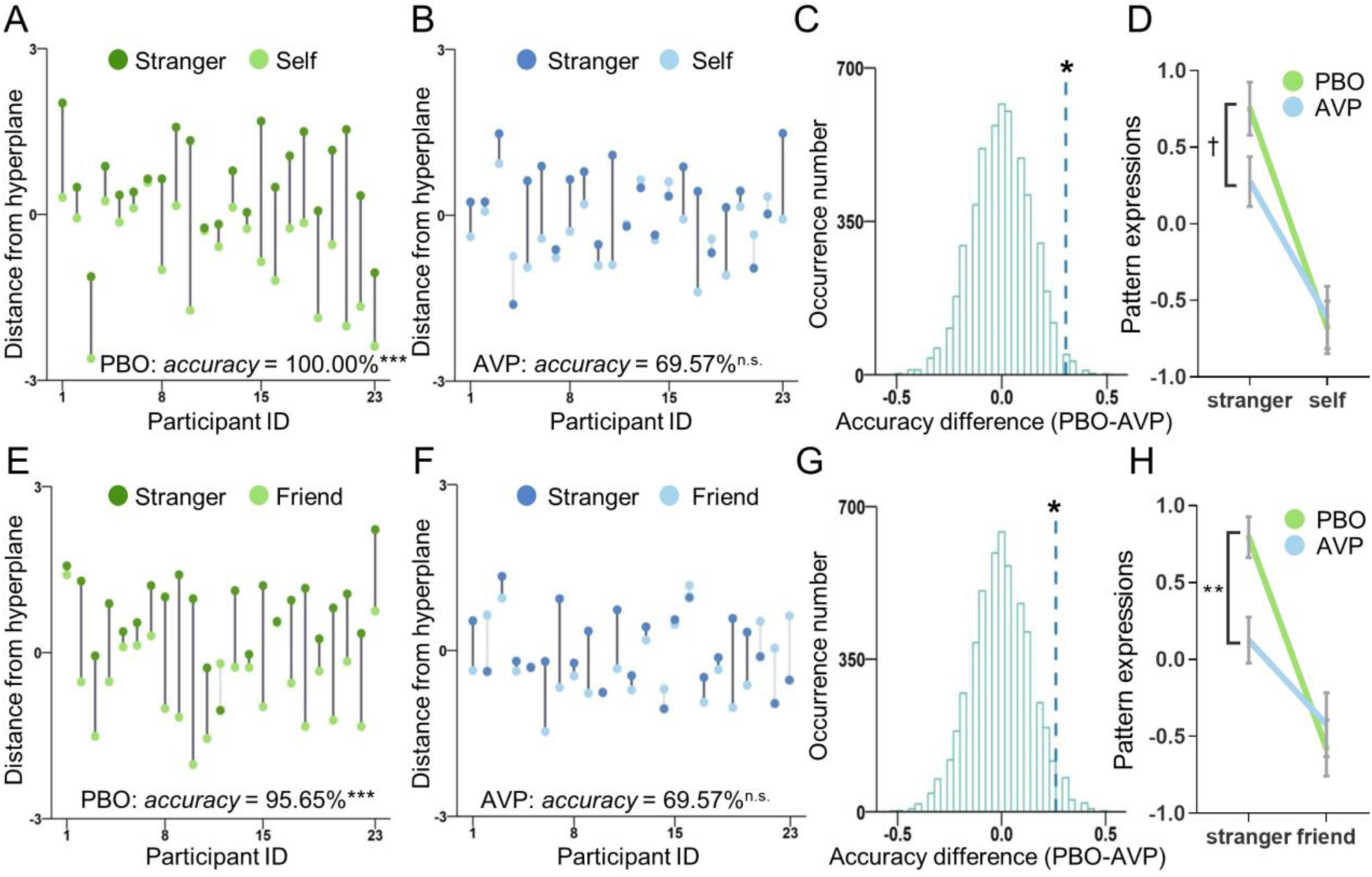
Effects of AVP on multivariate neural responses to the stranger compared to the self and friend. **(A)** Cross-validated distance from hyperplane derived from the stranger-self classifier across participants in the PBO treatment. Dark-gray lines indicate correct classification, and light-gray lines indicate incorrect classification. **(B)** Cross-validated distance from hyperplane derived from the stranger-self classifier across participants in the AVP treatment. **(C)** Permutation distribution of the differences in accuracy for the stranger-self classifier between PBO and AVP treatments. Blue dashed line indicates the value obtained from real score. **(D)** Pattern expressions for the stranger-self classifier as a function of target and treatment. **(E)** Cross-validated distance from hyperplane derived from the stranger-friend classifier across participants in the PBO treatment. **(F)** Cross-validated distance from hyperplane derived from the stranger-friend classifier across participants in the AVP treatment. **(G)** Permutation distribution of the differences in accuracy for the stranger-friend classifier between PBO and AVP treatments. **(H)** Pattern expressions the stranger-friend classifier as a function of target and treatment. PBO, placebo; AVP, arginine vasopressin; \*\*\**p* < 0.001; \*\**p* <0.01; \**p* < 0.05; ^†^*p* < 0.1; n.s., not significant.

The discrimination between friend and self was not significant under either PBO (*accuracy* = 69.57%, 95% *CI*: 50.76 ∼ 88.37%, *d* = 0.93, *p* = 9.31 × 10^-2^, **Fig. S5A**) or AVP treatment (*accuracy* = 65.22%, 95% *CI*: 45.75 ∼ 84.68%, *d* = 0.98, *p* = 2.10 × 10^-1^, **Fig. S5B**). Lastly, in the monetary outcome task, drug treatment had no significant effect on the discrimination between stranger and self, stranger and friend, or friend and self (**Table S4**).

The attenuating effects of AVP on neural distinctions between the stranger and the self or friend were further supported by two supplementary analyses. First, classification accuracy between the stranger and the combined self-friend condition was higher under the PBO treatment than under AVP (see supplementary results, **Fig. S6B∼S6F**). Second, cross-condition classifications between stranger-self and stranger-friend distinctions also performed better in the PBO treatment compared to AVP treatment (see supplementary results, **Fig. S7**).

Consistent with the results of classification performance, pattern expression analyses revealed that the target × treatment interaction was marginally significant for the stranger-self classifier (*F_1,44_* = 3.91, *p* = 5.42 × 10^-2^, *η*^2^ = 0.08, **Fig. 4D**), significant for the stranger-friend classifier (*F_1,44_* = 8.43, *p* = 5.76 × 10^-3^, *η*^2^ = 0.16, **Fig. 4H**) but not significant for the friend-self classifier (*F_1,44_* = 0.15, *p* = 6.98 × 10^-1^, *η*^2^*_p_* = 0.003). Follow-up simple effect analyses revealed higher pattern expression values for the stranger under the PBO treatment than AVP treatment in both the stranger-self (*mean difference ± s. e.* =0.48 ± 0.24, *p* = 5.12 × 10^-2^) and stranger-friend classifiers (*mean difference ± s. e.* = 0.67 ± 0.20, *p* = 1.74 × 10^-3^). However, there was no difference between drug treatments in the pattern expression values for self or friend (both *p* > 5.85 × 10^-1^). These findings suggest that AVP’s effects on differentiating the stranger from the self or the friend were primarily driven by its modulation of neural responses to the stranger.

### mPFC served as a robust contributing region to the classification between stranger and self or friend

The bootstrap test indicated that the mPFC, posterior cingulate cortex, thalamus, and midbrain robustly contributed to the discrimination between stranger and self (**Fig. 5A & Table S5**). Similarly, the mPFC and posterior cingulate cortex robustly contributed to the discrimination between stranger and friend (**Fig. 5B & Table S6**). These contributing regions were further validated using Haufe-transformed activation patterns (**Fig. S8**). Together with the univariate findings, these results provide converging evidence that AVP treatment modulated mPFC responses to the stranger, in comparison with responses to both self and friend.

**Fig. 5.**
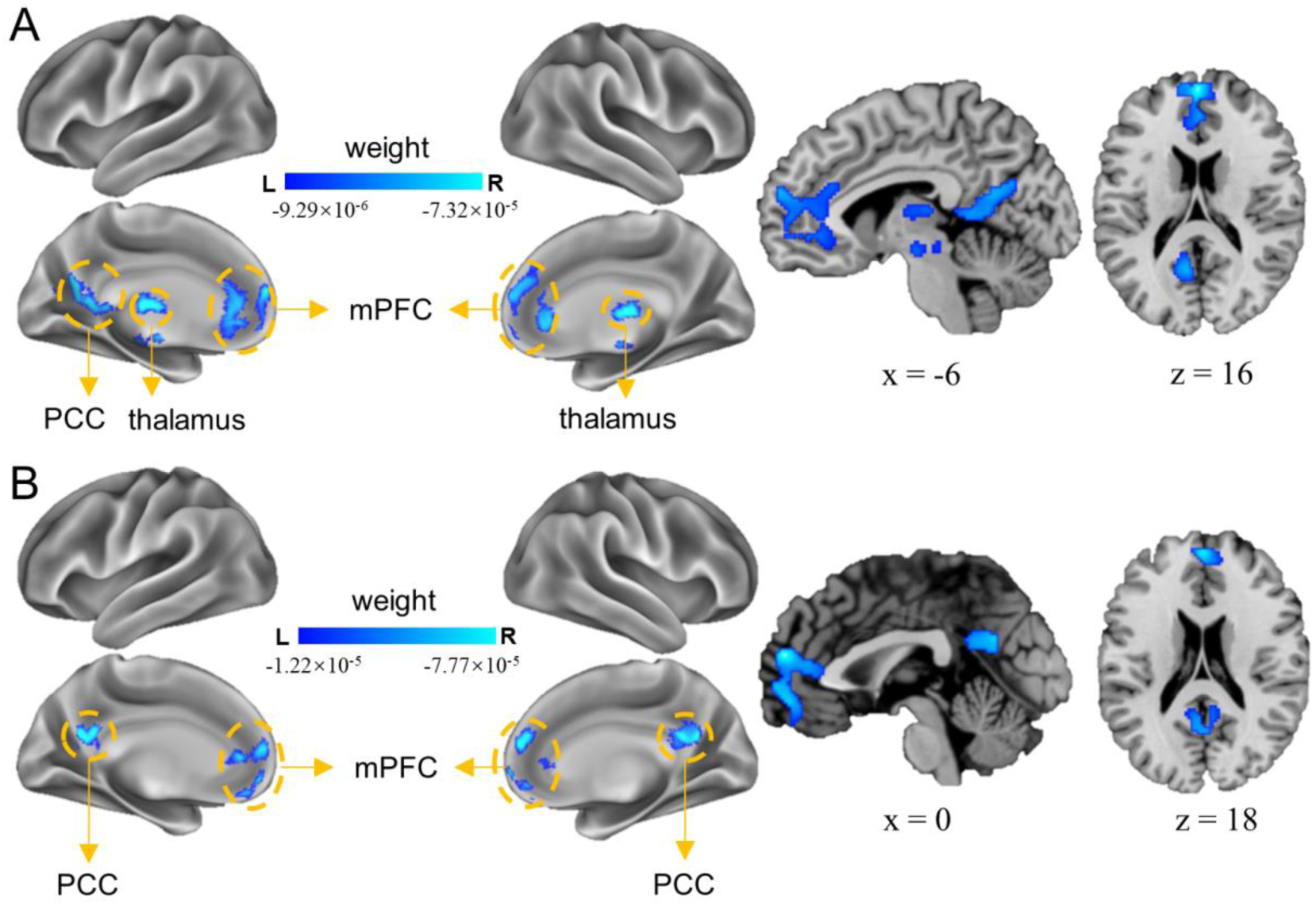
Brain regions robustly contributing to the distinction between stranger and self or friend. **(A)** The robust contributing regions in the stranger-self classifier. **(B)** The robust contributing regions in the stranger-friend classifier. mPFC, medial prefrontal cortex; PCC, posterior cingulate cortex; L, left; R, right.

### AVP reduced vmPFC connectivity with key nodes of the social cognitive network in response to strangers

The contrast analysis of [(stranger - self)_AVP_ vs. (stranger - self)_PBO_] using the vmPFC as a seed region revealed significant changes in connectivity with key nodes of the social cognitive network, including the temporoparietal junction (TPJ) and precuneus **(Fig. 6A & Table S7)**. Post-hoc comparisons indicated that AVP treatment diminished the functional connectivity between the vmPFC and these regions in response to the stranger, relative to both the self and friend (**Fig. 6B**). No significant changes in connectivity were found when using the dmPFC or rmPFC as seed regions.

**Fig. 6.**
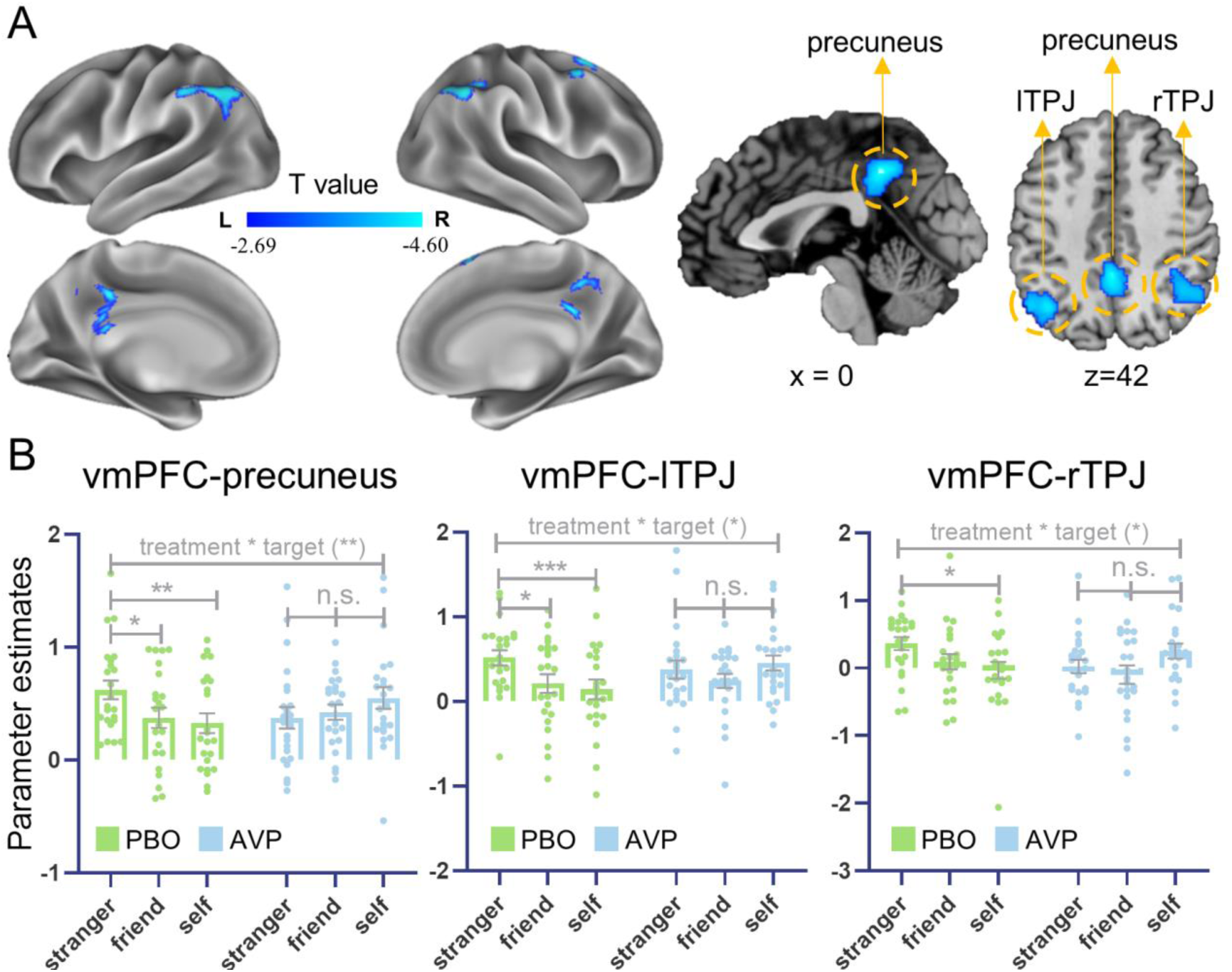
Effects of AVP on functional connectivity of vmPFC as a seed region. **(A)** Brain regions identified with the contrast of [(stranger - self)_AVP_ < (stranger - self)_PBO_]. (B) Post-hoc analyses for the identified precuneus and TPJ. vmPFC: ventromedial prefrontal cortex; lTPJ, left temporoparietal junction; rTPJ, right temporoparietal junction; PBO, placebo; AVP, arginine vasopressin; L, left; R, right. \*\*\**p* < 0.001, \*\**p* < 0.01, \**p* < 0.05, n.s. indicates not significant.

### Participants with AVP and PBO treatments were matched on sociodemographics, personality traits, and mood states

Participants with AVP and PBO treatments were well matched on sociodemographic measures, social preferences, personality traits and emotional states (for details, see **Table S8**).

## Discussion

This study examined the neural substrates underlying the effects of AVP on self-protective responses to spontaneous social comparison. Compared to placebo, AVP enhanced contrastive emotions to the social outcomes of a stranger, reducing satisfaction with positive evaluations and increasing satisfaction with negative evaluations of the stranger relative to those of the self or a friend. These results indicate that AVP heightened self-protective motives evoked by spontaneous social comparisons, an effect supported by converging evidence from changes in the mPFC activity, multivariate patterns, and functional connectivity. AVP increased mPFC activity to the stranger, reduced the neural distinctions between stranger and self/friend, and attenuated vmPFC connectivity with key nodes of the social cognitive network, including the TPJ and precuneus. The AVP modulation of the mPFC activity was particularly pronounced in socially dominant individuals. Notably, the effects of AVP on self-protective behavior and the mPFC function were specific to the social evaluation task. The current findings cannot be attributed to differences in personality traits or mood states between drug treatments, as these factors were well-matched across drug treatments. Together, these findings provide the first evidence that AVP promotes defensive responses to threats to the psychological self, revealing novel neuropsychological mechanisms underlying self-image protection.

Contrastive emotions like Schadenfreude and envy reflect subtle yet potent forms of defensive hostility that serve to protect desired self-views [23, 51]. These emotions are typically amplified by social comparative threats and predict subsequent defensive aggression [23]. Although the current study did not directly assess Schadenfreude and envy, our behavioral measure—derived from differences in satisfaction ratings in response to identical outcomes experienced by the stranger versus the self/friend—aligns well with the core unempathetic characteristic of contrastive emotions. That is, contrastive emotions are defined by affective incongruence across social targets, such that the same event is evaluated differently depending on whether it affects the self or another [25–26, 51]. AVP treatment increased this incongruence in contexts where participants were not explicitly asked to compare with others, highlighting its role in enhancing self-protective responses to threats arising from spontaneous social comparisons.

Our findings align with previous animal and human research showing that AVP lowers the threshold for interpreting social stimuli as threatening, and increases defensive hostility toward perceived threats [1, 3, 10–11]. However, this study is the first to demonstrate that AVP also promotes defense against threats to desired self-views, an inherently human challenge. These self-image threats are tied to the need to maintain a favorable self-image and social standing and entail detrimental consequences for well-being [16, 39]. For instance, longitudinal, experimental and meta-analytic studies consistently show that exposure to social comparative threats undermines well-being [20, 64–65]. These threats have become more prevalent and widespread in today’s era of social media [66], where individuals are frequently exposed to idealized portrayals of others. Therefore, it is adaptive for individuals to adopt strategies that protect or enhance their self-views when confronted with these challenges [18, 67]. Collectively, our findings provide novel evidence that a conserved function of AVP—defense against threats to survival or resources—has evolved to address contemporary social threats.

The effects of AVP on self-protective emotional responses were mediated by the mPFC function. Under placebo, mPFC responses to the stranger were lower than those to the self or a friend. In contrast, AVP treatment increased mPFC responses to the stranger, to levels comparable with those for the self and friend. Given the well-established role of the mPFC in encoding personal relevance of social targets [29], these findings suggest that AVP increases the perceived self-relevance of a potentially threatening other. Thus, our behavioral and neural findings reveal a seemingly paradoxical pattern: AVP enhanced unempathetic, distancing behavioral responses toward the stranger, yet reduced neural boundaries between the stranger and self/friend in the mPFC activity. This apparent divergence between behavioral and neural results reveals a novel mechanism whereby AVP promotes defensive threat responses—not by disengaging from threat, but by rendering the threatening other more self-relevant. Specifically, AVP appears to shift the representation of the stranger from a socially irrelevant figure under placebo to a psychologically salient competitor, thereby triggering the cascade of emotional and motivational processes underlying self-image defense.

This hypothesis is consistent with converging evidence from multiple domains. Threat-related stimuli are perceived as closer than neutral stimuli, a bias modulated by both the intensity of the threat and an individual’s sensitivity to social threats [68–69]. Likewise, when confronted with social comparative threats, individuals may seek proximity to the threat rather than to avoid it, echoing the adage “keep your friends close, but your enemies closer” [70]. At neural level, the mPFC not only encodes overlapping representations of the self and close others [30–31], but also plays a key role in tracking self-image threats and prompting defensive derogation and cognitive distortions [21, 34–35]. Our findings suggest that AVP may harness this circuitry to assimilate rivals into a psychologically proximate category, akin to close others in personal significance, thereby enhancing the motivational salience of social threats. This account is further corroborated by additional results from the current study.

First, AVP blunted the multivariate neural distinctions between the stranger and the self/friend. Under placebo, multivariate whole-brain patterns reliably distinguished the stranger from the self or friend, with the mPFC serving as a robust contributing region to the classification. However, AVP treatment reduced these distinctions to chance levels. Unlike the traditional univariate approach, which focuses on average neural activity within individual voxels, MVPA harnesses distributed patterns across multiple voxels, making it a more sensitive and rigorous method for detecting shared versus distinct neural representations [71–73]. Thus, our MVPA results offer compelling evidence that AVP assimilated mPFC responses to the stranger with those for the self and friend, strengthening the notion that AVP heightened the personal relevance of the stranger’s outcomes.

Second, the effects of AVP on mPFC activity were modulated by individual differences in dispositional dominance, with stronger effects in individuals higher in dominance. These findings indicate that AVP amplified the motivational significance of social comparative threats to a greater extent in dominant individuals. Consistent with our findings, socially dominant individuals exhibit an exaggerated sense of superiority and heightened sensitivity to social comparative threats, both of which drive stronger self-protective behaviors aimed at bolstering their social standing [41, 53]. For instance, dominant individuals are more inclined to seek proximity to social comparative threats rather than avoid them [70]. Moreover, animal studies have demonstrated that social dominance influences AVP and AVPR1a distribution and expression, leading to more aggressive responses to challenges in dominant individuals compared to submissive ones [43–44]. Collectively, these results highlight social dominance as a key modulator of AVP’s effects on threat processing and defensive aggression.

Third, AVP reduced the functional connectivity between the mPFC and key nodes of the social cognitive network, including the TPJ and precuneus, in response to the stranger. The TPJ is integral to distinguishing between self and others across various sociocognitive functions, potentially by enhancing task-relevant representations, suppressing non-relevant ones, or facilitating flexible switching between self- and other-related representations [74–75]. This region is particularly engaged by scenarios where self- and other-related representations conflict or when individuals must explicitly consider both representations [76]. Similarly, the precuneus plays a crucial role in maintaining clear boundaries between self and others [77]. It shows stronger engagement for third-person than first-person perspectives [78], contributes to shifting between these perspectives [79], and predicts better differentiation between self and others [80]. Therefore, the effects of AVP on the vmPFC-TPJ/precuneus connectivity provide additional evidence that AVP blurs the boundaries between stranger and self/friend, presumably by enhancing personal significance of the stranger.

Together, our findings suggest that AVP enhances self-protective emotional responses by reducing psychological distance to social threats and increasing their perceived immediacy. This is consistent with the social salience hypothesis, suggesting that AVP amplifies the relevance of social agents [81].

Several limitations warrant consideration. First, despite the collection of rich, multidimensional data, the sample size was relatively small for a between-subjects design. Replication in larger cohorts is necessary to assess the robustness and generalizability of the current findings. Second, the study focused exclusively on males, given AVP’s stronger and more consistent behavioral effects in men [82]. However, evidence also suggests AVP may promote defensive behaviors in females [10, 83], highlighting the need for future studies to examine sex-dependent effects. Third, although AVP is often linked to aggression, it can also promote affiliative strategies such as cooperation [45, 84]. Future work should explore whether AVP facilitates other forms of self-protection beyond defensive aggression. Fourth, behavioral effects of AVP were more pronounced in post-scan ratings than during-scan ratings, possibly due to the refined post-scan scale and/or timing-dependent effects of AVP [11]. These factors should be disentangled in future research. Fifth, contrastive emotional responses were indirectly measured as differential satisfaction ratings across targets to assess spontaneous social comparisons. Future studies should incorporate more direct measures of self-protective responses to social-comparative threats. Finally, although brain–behavior correlations and prior literature support our interpretations of identified regions, the fMRI findings primarily indicate regional involvement. Clarifying the specific functional roles of these regions will require studies using more targeted methodologies.

In summary, our study provides novel and converging evidence for the neuropsychological mechanisms underlying AVP’s role in self-protection against self-image threats prevalent in modern societies. Our findings indicate that the mPFC’s involvement in representing the personal relevance of social comparative threats mediates AVP’s effects, particularly among individuals with high social dominance orientation. These findings enhance our understanding of AVP’s role in human social behavior, provide new insights into the biological mechanisms of self-protection, and have important implications for the potential clinical application of AVP in psychiatric disorders characterized by self-protective deficits, such as depression and anxiety.

## Acknowledgements

This study was supported by the National Natural Science Foundation of China (32271126, 32020103008, 32371130), grant from Research Center for Brain Cognition and Human Development, Guangdong, China (No. 2024B0303390003), and grant from Striving for the First-Class, Improving Weak Links and Highlighting Features (SIH) Key Discipline for Psychology in South China Normal University.

## Conflict of interest

The authors are unaware of any conflicts of interest, financial or otherwise.

## Data and code availability

The behavioral data and the analysis code will be available at GitHub prior to publication. Due to the large size of the raw neuroimaging data, they will be available from the corresponding author upon request.

## Notes

### Competing Interest Statement

The authors have declared no competing interest.

