## Supplementary text, figures and tables for "Neural signature underlying the effect of intranasal vasopressin on emotional responses to spontaneous social comparison"

**Supplementary Material**

**Supplementary Methods**

**Tasks of the first visit**

*Medical history, friendship quality, sociodemographic information and personality traits*

Participants were asked to provide their medical histories, including histories of rhinitis, information about cold symptoms a week prior to the visit and information on drug intake two days prior to the visit. This information was used to screen participants for the drug experiment during the second visit.

Participants then rated the quality of their friendship with the friend who participated in the experiment with them. In particular, participants were asked about the duration of friendship (*How many months have you known the friend?*), frequency of contact (*How many times do you meet or communicate with this friend in each month?*), friendship quality (*How would you rate this friendship (range 1~10)?*), significance of the friendship (*How significant this friend is in your life (range 1~4)?*), and mutual consideration in decision (*How much does this friend consider your opinions when making important decisions (range 1~4)?, How much do you consider this friend’s opinions when making important decisions (range 1~4)?*).

Afterwards, participants were asked to fill in sociodemographic surveys and psychological questionnaires in order to control for differences in personality traits and emotional states between two groups receiving different drug treatments. Sociodemographic information included age, height, weight, number of siblings, relationship status, education level, birth sequence and family income. Participants also completed a series of psychological questionnaires that assessed their (i) preferences for social dominance with the social dominance orientation (SDO) scale [1], (ii) personality traits with the NEO five-factor inventory [2], (iii) empathy with the interpersonal reactivity index [3], (iv) alexithymia with the Toronto alexithymia scale-20 [4], (v) impulsiveness with the Barratt impulsiveness scale [5], (vi) self-esteem with the Rosenberg self-esteem scale [6], (vii) narcissism with the narcissistic personality inventory [7], (viii) social/trait anxiety with the fear of negative evaluation scale [8], interaction anxiousness scale [9], trait anxiety inventory [10], and self-rating anxiety scale [11], (ix) individualism and collectivism with the individualism and collectivism scale [12], (x) attachment style with the adult attachment scale [13], experiences in close relationships-relationship structures scale [14], and relationship scale questionnaire [15], (xi) childhood trauma with the childhood trauma questionnaire [16], (xii) father/mother love with the scales of father withdraw love and mother withdraw love [17], (xiii) intolerance of uncertainty with the intolerance of uncertainty scale [18], (xiv) autism with the autism quotient [19], (xv) depression with the Beck depression inventory [20] and self-rating depression [21], (xvi) aggression with the Buss-Perry aggression questionnaire [22], (xvii) interpersonal trust with the interpersonal trust scale [23], (xviii) loneliness with the UCLA loneliness scale [24], (xix) happiness with the depression-happiness scale [25], and (xx) reciprocity with the personal norm of reciprocity [26].

*Social preferences, political orientations, and self-introduction*

First, participants’ social preferences were measured with the following tasks: a social value orientation (SVO) task (Murphy et al., 2011; Murphy & Ackermann, 2014), an Ultimatum game (UG, (Güth et al., 1982)), a Dictator game (DG, (Forsythe et al., 1994; Kahneman et al., 1986)) and a Trust game (TG, (Berg et al., 1995)). In the SVO slider task, participants decided on the allocation of payoff between themselves and an anonymous other in 6 primary items and 9 secondary items. In UG, participants acted as the proposer to decide how to share a gain or a loss of 10 money units (MUs) with a responder. They were told that responders could either accept (both got paid accordingly) or reject (both got nothing in the gain frame or both lost 10 MUs in the loss frame) the offer. The rules of DG were similar with UG, except that the anonymous other could only accept the offers. In TG, participants started with an endowment of 15 MUs and decided whether to trust or not by sending any portion of the endowment to a trustee and to keep the remainder of the endowment. The shared money was then tripled in value and passed on to the trustee, who would decide how much to return. Participants played 10 rounds of one-shot UG, DG, and TG, and their preferences in each game were calculated by averaging decisions across rounds.

Second, participants’ political orientations were measured by answering questions about the 2016 United States (U.S.) presidential election, which was ongoing when the current experiment was conducted. Participants were asked which candidate they preferred to be the next president of US (Donald Trump or Hillary Clinton).

Third, participants were asked to deliver a self-introduction, lasting a minimum of 2 minutes, which was captured on video. They were instructed to discuss their strengths, weaknesses, hobbies, and any other aspects relevant to themselves.

After the self-introduction, participants were asked if they agreed to share their personal profiles—including sociodemographic information, personality traits, social preferences, attitudes, and self-introduction—to others for a follow-up impression forming and social evaluation experiment. Specifically, participants were told that eight evaluators (four males and four females) would form impressions of them based on their personal profiles and evaluate them with personality trait adjectives. All participants agreed to proceed and returned for the social evaluation/monetary outcome task in the fMRI scanner about one week later. Unbeknownst to participants, their personal profiles were not shared with evaluators and the experiment procedure was set to convince participants that the social evaluations they received during the second visit were based on their personal profiles.

**Supplementary Results**

**Satisfaction ratings in the monetary outcome task**

In the monetary outcome task, LMMs of during-scan and post-scan satisfaction ratings revealed a significant 2-way interaction of treatment × outcome (during-scan ratings: *χ^2^_1_* = 173.71, *p* = 1.15 × 10^-39^, **Fig. S2A**; post-scan ratings: *χ^2^_1_* = 224.11, *p* = 1.15 × 10^-50^, **Fig. S2B**). Follow-up simple effect analyses indicated that satisfaction ratings of monetary gains were lower under AVP treatment compared to PBO treatment (during-scan: *parameter estimates* ± *s. e.* = -0.21 ± 0.05, *p* = 1.59 × 10^-4^; post-scan: *parameter estimates* ± *s. e.* = -0.38 ± 0.10, *p* = 1.82 × 10^-4^). In contrast, satisfaction ratings of monetary losses were higher under AVP treatment compared to PBO treatment (during-scan: *parameter estimates* ± *s. e.* = 0.27 ± 0.05, *p* = 2.66 × 10^-6^; post-scan: *parameter estimates* ± *s. e.* = 0.61 ± 0.10, *p* = 3.87 × 10^-8^). Other effects associated with treatment were not significant (**Table S3**).

**Exploratory whole-brain univariate analyses**

The exploratory univariate whole-brain analyses revealed no significant effects of drug treatment in the social evaluation task. For the monetary outcome task, the interaction between treatment and target revealed activation in the middle cingulate cortex/supplementary motor area (**Fig. S3**).

**The influence of dispositional dominance on AVP-induced changes in dmPFC and vmPFC activity**

The interaction between SDO score and treatment was not significant for the activity of dmPFC (stranger vs. self: *F_1, 40_* = 0.52, *p* = 4.73 × 10^-1^, **Fig. S4A**; stranger vs. friend: *F_1, 40_* = 3.47, *p* = 6.99 × 10^-2^, **Fig. S4B**) or vmPFC (stranger vs. self: *F_1, 40_* = 2.08, *p* = 1.57 × 10^-1^, **Fig. S4C**; stranger vs. friend: *F_1, 40_* = 1.74, *p* = 1.95 × 10^-1^, **Fig. S4D**). However, post-hoc analyses of simple slopes revealed similar patterns with rmPFC activity. Under AVP treatment, higher SDO scores were positively associated with increased dmPFC and vmPFC responses to the stranger relative to both self (dmPFC: *β* ± *s. e.* = 0.02 ± 0.03, *p* = 5.46 × 10^-1^, **Fig. S4A**; vmPFC: *β* ± *s. e.* = 0.06 ± 0.03, *p* = 2.60 × 10^-2^, **Fig. S4C**) and friend (dmPFC: *β* ± *s. e.* = 0.07 ± 0.03, *p* = 1.54 × 10^-2^, **Fig. S4B**; vmPFC: *β* ± *s. e.* = 0.08 ± 0.04, *p* = 3.93 × 10^-2^, **Fig. S4D**). In contrast, under PBO treatment, SDO scores were not significantly associated with dmPFC/vmPFC activity for either contrast of stranger versus self (dmPFC: *β* ± *s. e.* = -0.03 ± 0.05, *p* = 6.21 × 10^-1^, **Fig. S4A**; vmPFC: *β* ± *s. e.* = 0.02 ± 0.05, *p* = 6.77 × 10^-1^, **Fig. S4C**) or stranger versus friend (dmPFC: *β* ± *s. e.* = -0.04 ± 0.05, *p* = 4.41 × 10^-1^, **Fig. S4B**; vmPFC: *β* ± *s. e.* = -0.03 ± 0.07, *p* = 7.10 × 10^-1^, **Fig. S4D**).

**Effects of AVP on the contrast between stranger and the combined self-friend condition**

For the moderation effect on the rmPFC activity, the complementary analyses with the combined self-friend condition revealed a significant interaction between SDO scores and treatment (*F_1, 40_* = 4.94, *p* = 3.20 × 10^-2^, **Fig. S6A**). Simple slope analyses indicated that higher SDO scores were positively associated with increased rmPFC responses to the stranger relative to the self-friend condition under AVP treatment (*β* ± *s. e.* = 0.10 ± 0.03, *p* = 2.05 × 10^-3^) but not under PBO treatment (*β* ± *s. e.* = -0.04 ± 0.06, *p* = 4.37 × 10^-1^).

For the MVPA results, the classifier discriminating stranger vs. the combined self-friend condition demonstrated high accuracy in the PBO treatment (*accuracy* = 95.65%, 95% *CI*: 87.32 ~ 100.00%, *d* = 2.29, *p* = 5.72 × 10^-6^, **Fig. S6B**) but not in the AVP treatment (*accuracy* = 60.87%, 95% *CI*: 40.92 ~ 80.82%, *d* = 0.49, *p* = 4.05 × 10^-1^, **Fig. S6C**). Permutation tests revealed a significant difference in discrimination accuracy between PBO and AVP treatments (*p* = 7.20 × 10^-3^, **Fig. S6D**). These classification results were further supported by the analyses of pattern expressions derived from the stranger vs. self-friend classifier, which revealed a significant target × treatment interaction (*F_1,44_* = 7.25, *p* = 1.00 × 10^-2^, *η*^2^*_p_* = 0.14, **Fig. S6E**). Follow-up simple effect analyses revealed higher pattern expression values for the stranger under the PBO treatment than AVP treatment (*mean difference* ± *s. e.* = 0.64 ± 0.23, *p* = 7.01 × 10^-3^), while there was no difference between drug treatments in the pattern expression values for the combined self-friend condition (*mean difference* ± *s. e.* = -0.16 ± 0.29, *p* = 5.84 × 10^-1^).

Lastly, the bootstrap test on the weights of stranger vs. self-friend classifier identified similar robust contributing regions with those found in the separate classifiers, including the mPFC, posterior cingulate cortex, and thalamus (**Fig. S6F**).

**AVP diminished the cross-condition classifications between stranger-self and stranger-friend distinctions**

The cross-condition classification was demonstrated under the PBO treatment (the stranger-self classifier tested on the stranger-friend distinction: *accuracy* = 82.61%, 95% *CI*: 67.12 ~ 98.10%, *d* = 1.80, *p* = 2.60 × 10^-3^, **Fig. S7A**; the stranger-friend classifier tested on the stranger-self distinction: *accuracy* = 100.00%, 95% *CI*: 100.00 ~ 100.00%, *d* = 2.23, *p* = 2.38 × 10^-7^, **Fig. S7D**). In contrast, AVP treatment reduced the cross-condition classification to the chance level (the stranger-self classifier tested on the stranger-friend distinction: *accuracy* = 60.87%, 95% *CI*: 40.92 ~ 80.82%, *d* = 0.25, *p* = 4.05 × 10^-1^, **Fig. S7B**; the stranger-friend classifier tested on the stranger-self distinction: *accuracy* = 52.17%, 95% *CI*: 31.76 ~ 72.59%, *d* = 0.26, *p* = 1.00, **Fig. S7E**). The classification accuracy was higher for the PBO treatment than the AVP treatment (the stranger-self classifier tested on the stranger-friend distinction: *p* = 3.42 × 10^-2^, **Fig. S7C**; the stranger-friend classifier tested on the stranger-self distinction: *p* < 2.00 × 10^-4^, **Fig. S7F**).

**Supplementary Figures and Tables**

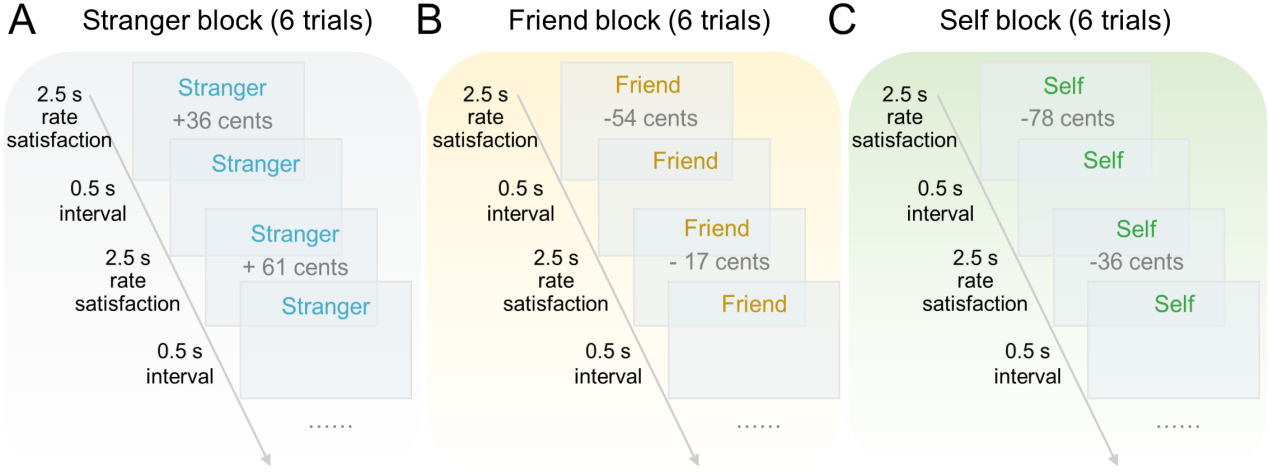

**Fig. S1 | The timeline of the monetary outcome task. (A)** Timeline of the monetary outcome task for a stranger block. **(B)** Timeline of the monetary outcome task for a friend block. **(C)** Timeline of the monetary outcome task for a self block. Each fMRI run consisted of separate blocks corresponding to six experimental conditions defined by the combination of three targets (stranger, friend, self) and two evaluation types (positive, negative). Each block began with a 3-second cue indicating the current target (“stranger,” “friend,” or “self”), followed by six consecutive evaluations directed to that target. Each evaluation was presented for 2-second followed by 0.5-second of blank screen. Two successive blocks were separated by a 10-second rest period with a central fixation.

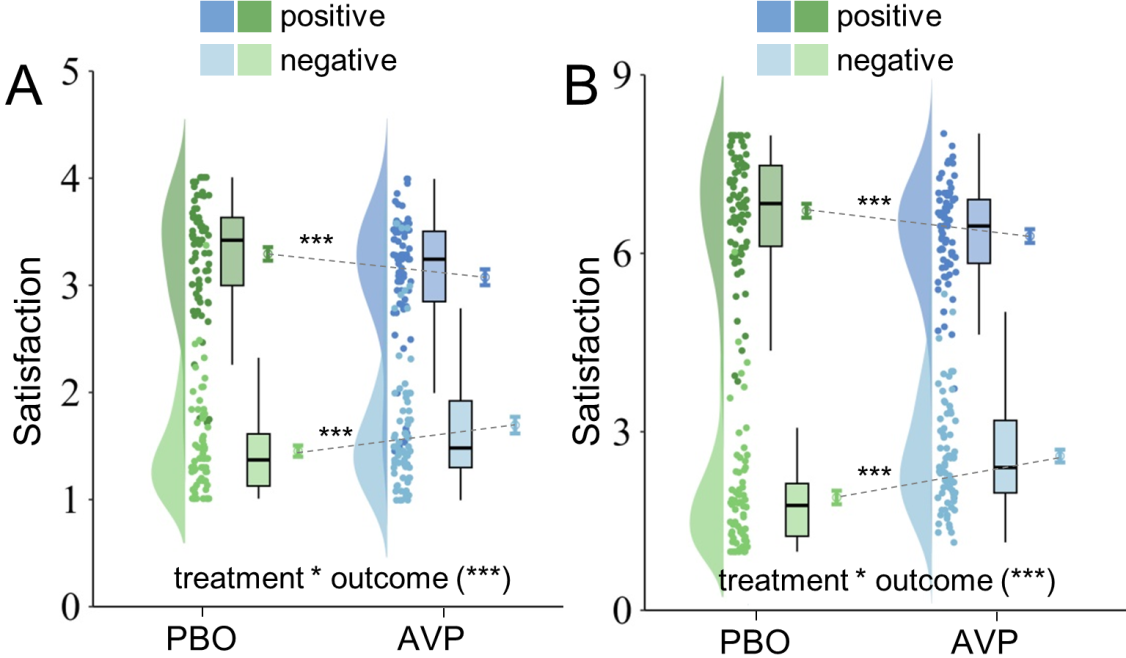

**Fig. S2 | Results of satisfaction ratings in the monetary outcome task.** A significant 2-way interaction of treatment × outcome was found in both **(A)** during-scan and **(B)** post-scan satisfaction ratings. PBO, placebo; AVP, arginine vasopressin. Error bars show standard error. ****p* < 0.001.

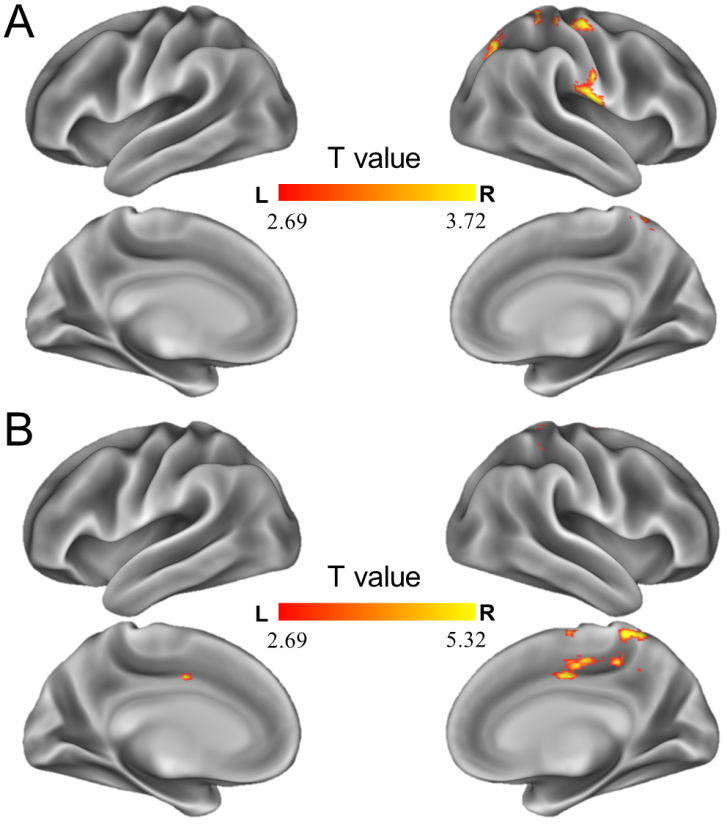

**Fig. S3 | Exploratory whole-brain analysis on the interaction between drug and treatment for both tasks.** **(A)** The interaction did not reveal significant brain activations for social evaluation task. Brain regions identified with *p* < 0.005 and cluster size > 100 voxels were illustrated. **(B)** The interaction revealed middle cingulate cortex/supplementary motor area (peak MNI coordinates: x, y, z = 14, -24, 54 mm, cluster size = 938 voxels) for monetary outcome task. L, left; R, right.

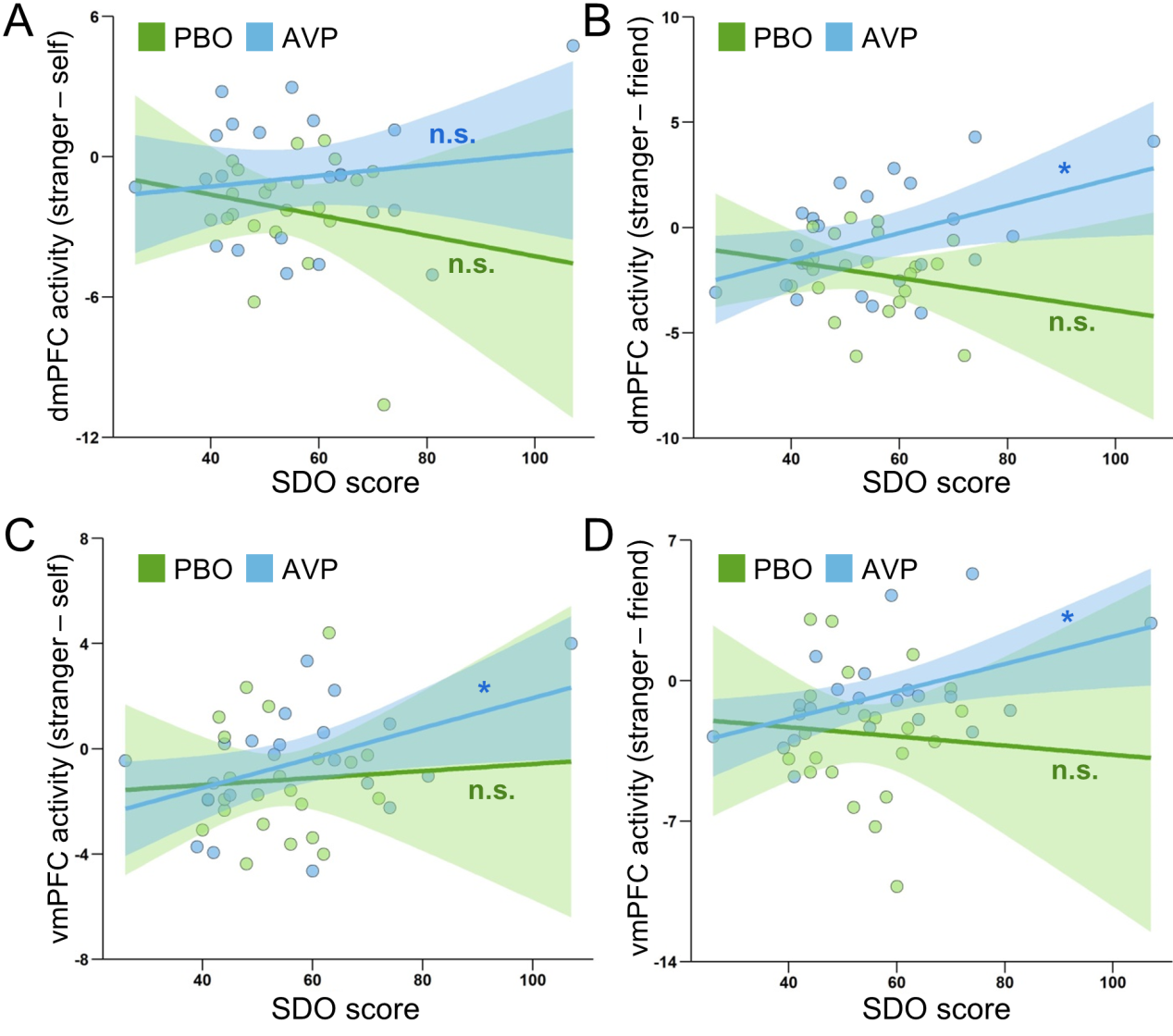

**Fig. S4 | Modulations of dispositional dominance on the AVP-induced changes in dmPFC and vmPFC activity. (A)** Associations between SDO scores and dmPFC activity to the contrast of stranger vs. self as a function of treatment. **(B)** Associations between SDO scores and dmPFC activity to the contrast of stranger vs. friend as a function of treatment. **(C)** Associations between SDO scores and vmPFC activity to the contrast of stranger vs. self as a function of treatment. **(D)** Associations between SDO scores and vmPFC activity to the contrast of stranger vs. friend as a function of treatment. Lines represent fitted correlations, with shaded areas indicating 95% confidence intervals. dmPFC, dorsomedial prefrontal cortex; vmPFC, ventromedial prefrontal cortex; PBO, placebo; AVP, arginine vasopressin; SDO, social dominance orientation. **p* < 0.05; n.s., not significant.

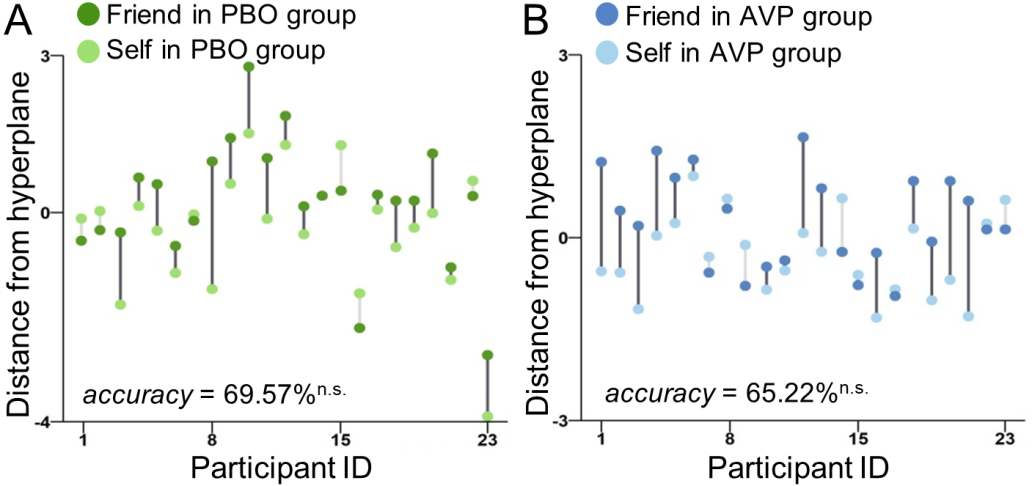

**Fig. S5 | Effects of AVP on multivariate neural distinction between friend and self. (A)** Cross-validated distance from hyperplane derived from the friend-self classifier across participants in the PBO treatment. Dark-gray lines indicate correct classification, and light-gray lines indicate incorrect classification. **(B)** Cross-validated distance from hyperplane derived from the friend-self classifier across participants in the AVP treatment. PBO, placebo; AVP, arginine vasopressin; n.s., not significant.

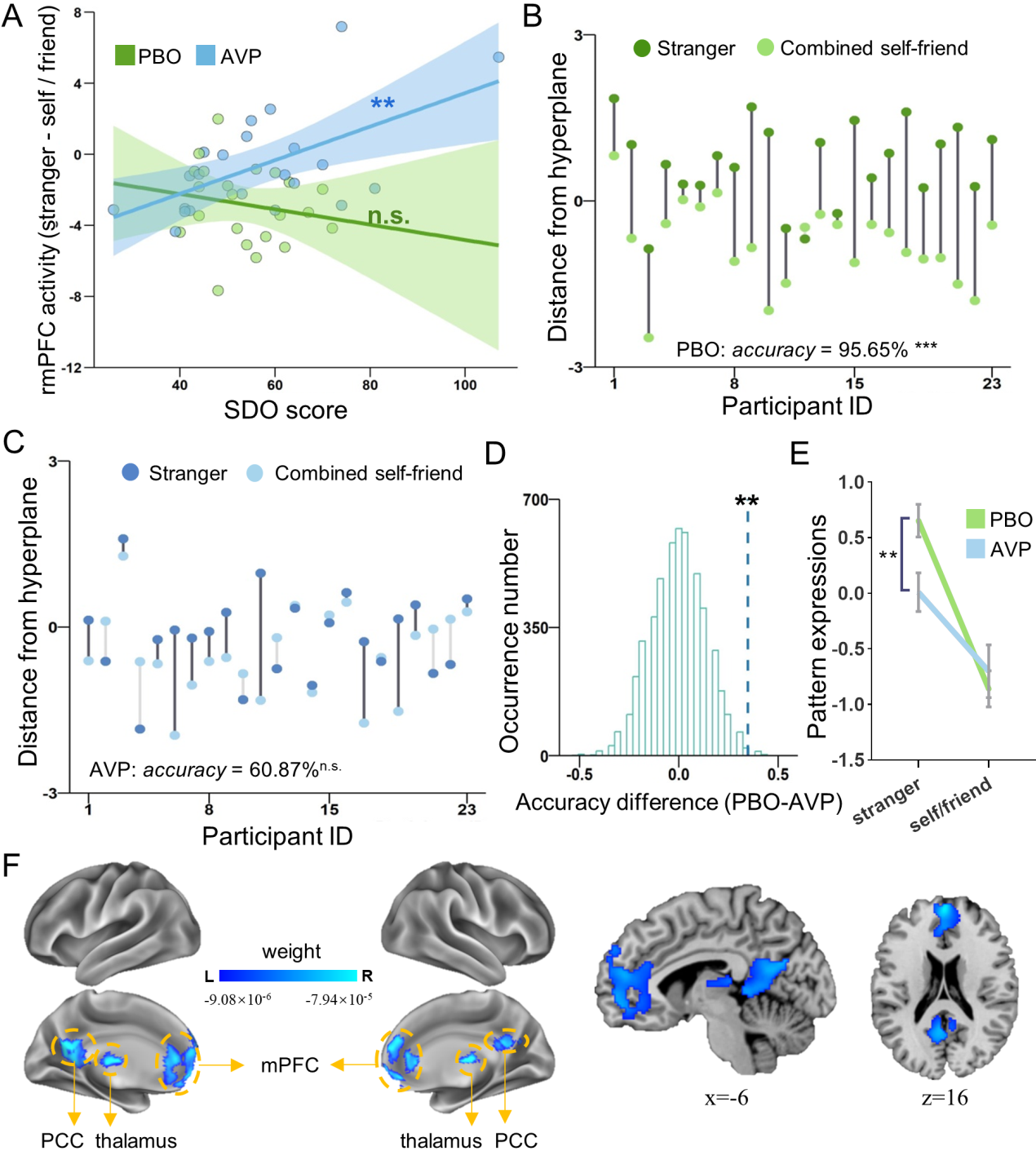

**Fig. S6 | Complementary results with the combined self-friend condition.** **(A)** Associations between SDO scores and rmPFC activity to the contrast of stranger vs. combined self-friend as a function of treatment. Lines represent fitted correlations, with shaded areas indicating 95% confidence intervals. **(B)** Cross-validated distance from hyperplane derived from the stranger vs. combined self-friend classifier across participants in the PBO treatment. Dark-gray lines indicate correct classification, and light-gray lines indicate incorrect classification. **(C)** Cross-validated distance from hyperplane derived from the stranger vs. combined self-friend classifier across participants in the AVP treatment. **(D)** Permutation distribution of the differences in accuracy for the stranger vs. combined self-friend classifier between PBO and AVP treatments. Blue dashed line indicates the value obtained from real scores. **(E)** Pattern expressions for the stranger vs. combined self-friend classifier as a function of target and treatment. Error bars show standard error. **(F)** The robust contributing regions in the stranger vs. combined self-friend classifier. rmPFC, rostromedial prefrontal cortex; mPFC, medial prefrontal cortex; PCC, posterior cingulate cortex; PBO, placebo; AVP, arginine vasopressin; SDO, social dominance orientation; L, left; R, right; ****p* < 0.001; ***p* < 0.01; n.s., not significant.

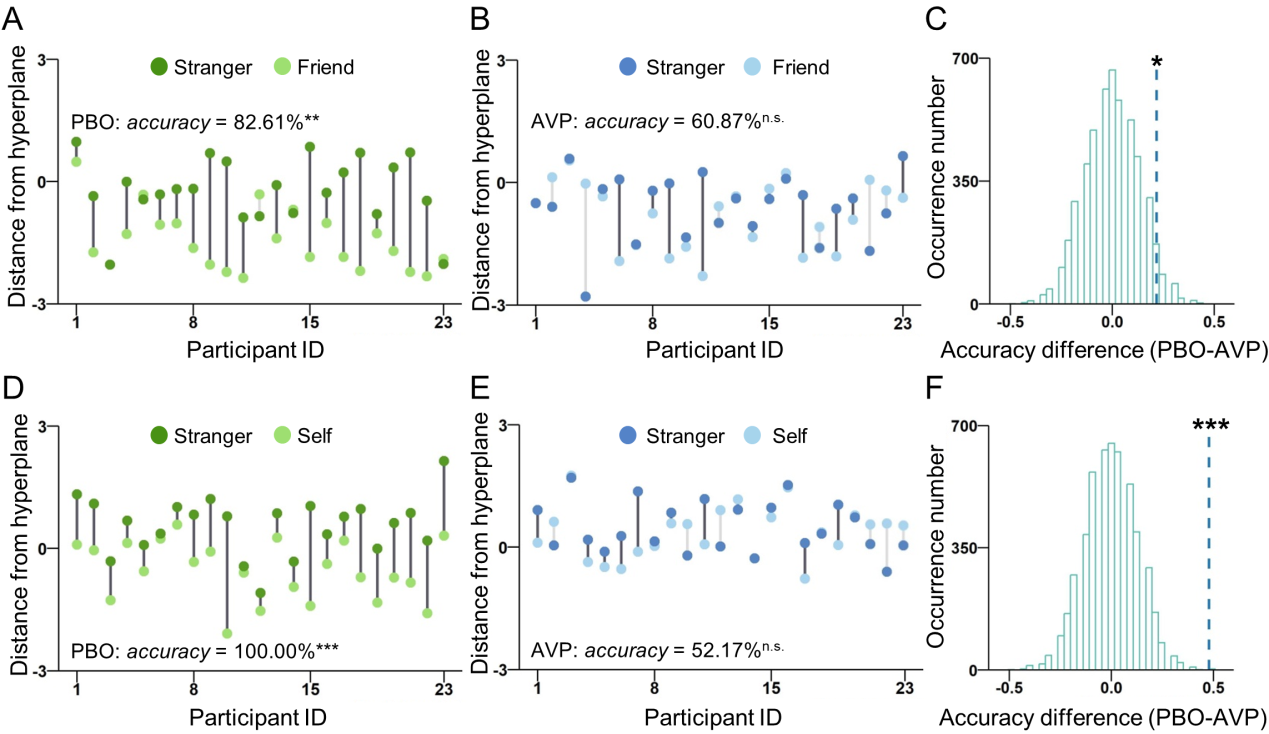

**Fig. S7 | Results of cross-condition classification. (A)** Cross-validated distance from hyperplane for the stranger and friend derived from the stranger-self classifier across participants in the PBO treatment. Dark-gray lines indicate correct classification, and light-gray lines indicate incorrect classification. **(B)** Cross-validated distance from hyperplane for the stranger and friend derived from the stranger-self classifier across participants in the AVP treatment. **(C)** Permutation distribution of the differences between PBO and AVP treatments in accuracy for the stranger vs. friend distinction based on the stranger-self classifier. Blue dashed line indicates the value obtained from real scores. **(D)** Cross-validated distance from hyperplane for the stranger and self derived from the stranger-friend classifier across participants in the PBO treatment. **(E)** Cross-validated distance from hyperplane for the stranger and self derived from the stranger-friend classifier across participants in the AVP treatment. **(F)** Permutation distribution of the differences between PBO and AVP treatments in accuracy for the stranger vs. self distinction based on the stranger-friend classifier. PBO, placebo; AVP, arginine vasopressin; ****p* < 0.001; ***p* < 0.01; **p* < 0.05; n.s., not significant.

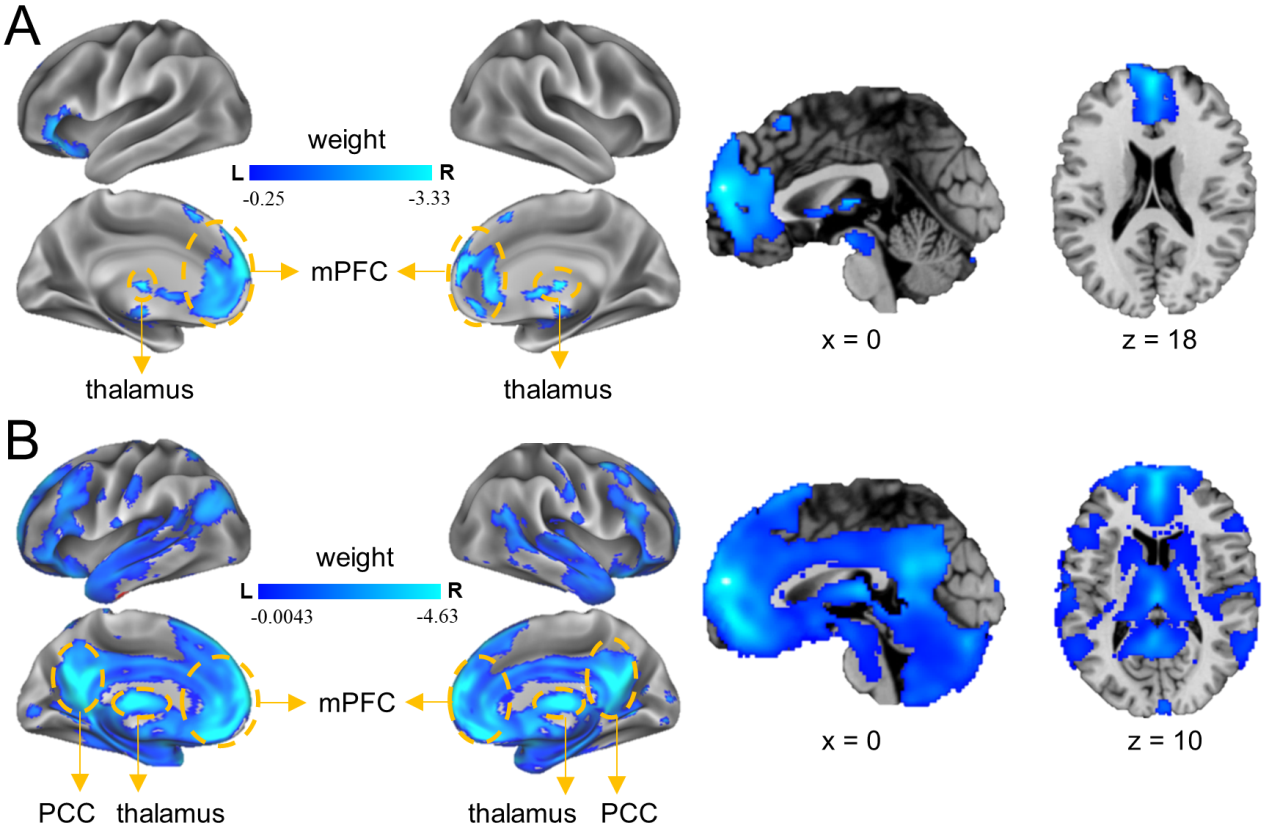

**Fig. S8 | Brain regions robustly contributing to the distinction between stranger and self or friend identified with Haufe-transformed activation patterns. (A)** The robust contributing regions in the stranger-self classifier. **(B)** The robust contributing regions in the stranger-friend classifier. mPFC, medial prefrontal cortex; PCC, posterior cingulate cortex; L, left; R, right.

**Table S1 | Items used in the social evaluation task.**

| **Positive attributes** | **Negative attributes** |
| --- | --- |
| generous (慷慨) | rude (粗鲁) |
| virtuous (厚道) | superficial (肤浅) |
| warmhearted (热心) | fickle (浮躁) |
| pragmatic (务实) | odd (古怪) |
| sensitive (敏锐) | stubborn (固执) |
| efficient (干练) | void (空虚) |
| surefooted (踏实) | detached (冷漠) |
| careful (细心) | stingy (吝啬) |
| clement (宽厚) | reckless (鲁莽) |
| rigorous (严谨) | outdated (落伍) |
| humane (仁爱) | numb (麻木) |
| composed (沉着) | blind (盲目) |
| positive (积极) | cowardly (懦弱) |
| steady (稳重) | shallow (浅薄) |
| honest (诚实) | rigid (死板) |
| diligent (努力) | bored (无聊) |
| studious (好学) | ignorant (无知) |
| independent (独立) | narrow (狭隘) |
| upright (正直) | vain (虚荣) |
| responsible (负责) | vulgar (庸俗) |
| self-supporting (自立) | stupid (愚昧) |
| strong (坚强的) | obtrusive (张扬) |
| kind (善良) | self-abased (自卑) |
| sincere (真诚) | artificial (做作) |

All of the items were presented in Chinese in the experiment as shown in the brackets.

**Table S2 | All statistical effects of during-scan and post-scan satisfaction ratings in the social evaluation and monetary outcome tasks**

|  | Effects | Likelihood ratio | *p* |
| --- | --- | --- | --- |
| social evaluation task | |  |  |
| during-scan ratings | treatment | χ^2^(1) = 7.49 | .006 |
|  | outcome | χ^2^(1) = 191.50 | <.001 |
|  | target | χ^2^(2) = 38.78 | <.001 |
|  | treatment × outcome | χ^2^(1) = 58.76 | <.001 |
|  | treatment × target | χ^2^(2) = 16.88 | <.001 |
|  | outcome × target | χ^2^(2) = 47.71 | <.001 |
|  | treatment × outcome × target | χ^2^(2) = 5.44 | .066 |
| post-scan ratings | treatment | χ^2^(1) = 0.91 | .339 |
|  | outcome | χ^2^(1) = 223.03 | <.001 |
|  | target | χ^2^(2) = 6.00 | .050 |
|  | treatment × outcome | χ^2^(1) = 129.90 | <.001 |
|  | treatment × target | χ^2^(2) = 2.14 | .342 |
|  | outcome × target | χ^2^(2) = 199.92 | <.001 |
|  | treatment × outcome × target | χ^2^(2) = 34.18 | <.001 |
| monetary outcome task | |  |  |
| during-scan ratings | treatment | χ^2^(1) = 0.39 | .530 |
|  | outcome | χ^2^(1) = 375.57 | <.001 |
|  | target | χ^2^(2) = 1.99 | .369 |
|  | treatment × outcome | χ^2^(1) = 173.71 | <.001 |
|  | treatment × target | χ^2^(2) = 1.31 | .519 |
|  | outcome × target | χ^2^(2) = 293.27 | <.001 |
|  | treatment × outcome × target | χ^2^(2) = 1.79 | .408 |
| post-scan ratings | treatment | χ^2^(1) = 1.62 | .203 |
|  | outcome | χ^2^(1) = 546.69 | <.001 |
|  | target | χ^2^(2) = 13.13 | .001 |
|  | treatment × outcome | χ^2^(1) = 224.11 | <.001 |
|  | treatment × target | χ^2^(2) = 5.02 | .081 |
|  | outcome × target | χ^2^(2) = 330.85 | <.001 |
|  | treatment × outcome × target | χ^2^(2) = 3.99 | .136 |

*Note:* The *p* values of LMM were obtained through likelihood ratio tests comparing the full model, which included the effect of interest, against a reduced model without it [27-28].

**Table S3 | All statistical effects on the activity of mPFC subregions.**

|  | Effect | *F* | *p* | *η^2^_p_* |
| --- | --- | --- | --- | --- |
| social evaluation task | |  |  |  |
| dmPFC | treatment | 0.003 | .959 | .000 |
|  | outcome | 5.541 | .023 | .112 |
|  | target | 11.135 | <.001 | .202 |
|  | treatment × outcome | 0.005 | .946 | .000 |
|  | treatment × target | 3.390 | .038 | .072 |
|  | outcome × target | 1.538 | .222 | .034 |
|  | treatment × outcome × target | 1.965 | .150 | .043 |
| rmPFC | treatment | 1.706 | .198 | .037 |
|  | outcome | 0.238 | .628 | .005 |
|  | target | 10.068 | <.001 | .186 |
|  | treatment × outcome | 1.264 | .267 | .028 |
|  | treatment × target | 3.593 | .035 | .075 |
|  | outcome × target | 1.468 | .237 | .032 |
|  | treatment × outcome × target | 0.604 | .530 | .014 |
| vmPFC | treatment | 0.179 | .674 | .004 |
|  | outcome | 0.003 | .955 | .000 |
|  | target | 12.007 | <.001 | .214 |
|  | treatment × outcome | 2.930 | .094 | .062 |
|  | treatment × target | 4.457 | .017 | .092 |
|  | outcome × target | 1.010 | .368 | .022 |
|  | treatment × outcome × target | 0.432 | .648 | .010 |
| monetary outcome task | |  |  |  |
| dmPFC | treatment | 0.422 | .519 | .009 |
|  | outcome | 0.469 | .497 | .011 |
|  | target | 1.597 | .209 | .035 |
|  | treatment × outcome | 0.068 | .795 | .002 |
|  | treatment × target | 0.995 | .371 | .022 |
|  | outcome × target | 0.357 | .687 | .008 |
|  | treatment × outcome × target | 0.037 | .957 | .001 |
| rmPFC | treatment | 0.892 | .350 | .020 |
|  | outcome | 1.000 | .323 | .022 |
|  | target | 0.441 | .621 | .010 |
|  | treatment × outcome | 0.058 | .810 | .001 |
|  | treatment × target | 2.068 | .139 | .045 |
|  | outcome × target | 0.033 | .964 | .001 |
|  | treatment × outcome × target | 3.047 | .054 | .065 |
| vmPFC | treatment | 0.148 | .702 | .003 |
|  | outcome | 0.579 | .451 | .013 |
|  | target | 1.247 | .289 | .028 |
|  | treatment × outcome | 0.185 | .669 | .004 |
|  | treatment × target | 1.620 | .208 | .036 |
|  | outcome × target | 0.567 | .568 | .013 |
|  | treatment × outcome × target | 1.745 | .181 | .038 |

dmPFC, dorsomedial prefrontal cortex; rmPFC, rostromedial prefrontal cortex; vmPFC, ventromedial prefrontal cortex.

**Table S4 | Classification performance of each target pair based on whole-brain neural patterns for the monetary outcome task.**

|  | PBO group | AVP group |
| --- | --- | --- |
| stranger vs. self | 39.13% [95% *CI*: 19.18 ~ 59.08%] | 69.57% [95% *CI*: 50.76 ~ 88.37%] |
| stranger vs. friend | 65.22% [95% *CI*: 45.75 ~ 84.68%] | 56.52% [95% *CI*: 36.26 ~ 76.78%] |
| friend vs. self | 69.57% [95% *CI*: 50.76 ~ 88.37%] | 65.22% [95% *CI*: 45.75 ~ 84.68%] |
| stranger vs. combined self-friend | 47.83% [95% *CI*: 27.41 ~ 68.24%] | 56.52% [95% *CI*: 36.26 ~ 76.78%] |

All the classifiers above performed at chance level. PBO, placebo; AVP, arginine vasopressin; CI, confidence interval.

**Table S5 | Brain regions robustly contributing to the distinction between stranger and self**

| Brain Regions | Cluster size (voxel) | L/R | MNI Coordinates of Local Maxima (mm) | | | Weight |
| --- | --- | --- | --- | --- | --- | --- |
|  |  |  | x | y | z |  |
| Medial prefrontal cortex | 1380 | L/R | 0 | 58 | 18 | -7.32 × 10^-5^ |
| Posterior cingulate cortex | 284 | L/R | -4 | -42 | 4 | -5.34 × 10^-5^ |
| Midbrain | 203 | L/R | -2 | -14 | -14 | -3.77 × 10^-5^ |
| Thalamus | 201 | L/R | 0 | -22 | 10 | -6.45 × 10^-5^ |

L, left; R, right.

**Table S6 | Brain regions robustly contributing to the distinction between stranger and friend**

| Brain Regions | Cluster size (voxel) | L/R | MNI Coordinates of Local Maxima (mm) | | | Weight |
| --- | --- | --- | --- | --- | --- | --- |
|  |  |  | x | y | z |  |
| Medial prefrontal cortex | 766 | L/R | 0 | 54 | 16 | -7.77 × 10^-5^ |
| Posterior cingulate cortex | 397 | L/R | 0 | -62 | 26 | -6.38 × 10^-5^ |

L, left; R, right.

**Table S7 | Brain regions exhibiting AVP-induced changes in functional connectivity with ventromedial prefrontal cortex**

| Brain Regions | Cluster size | L/R | MNI Coordinates of Local Maxima (mm) | | | Local Maxima |
| --- | --- | --- | --- | --- | --- | --- |
|  | (voxel) |  | x | y | z | T |
| precuneus | 429 | L/R | 0 | -44 | 40 | -4.60 |
| temporoparietal junction | 399 | L | -38 | -64 | 40 | -4.18 |
| temporoparietal junction | 430 | R | 52 | -56 | 46 | -3.84 |
| supplementary motor area | 363 | R | 12 | 18 | 62 | -4.13 |

L, left; R, right.

**Table S8 | Sociodemographic information, psychological questionnaires and mood measurements.**

|  | PBO group (n = 24) | AVP group （n = 24) |  |  |
| --- | --- | --- | --- | --- |
|  | Percentage | Percentage | Person  Chi-square | *p* |
| marital status (single/in a relationship) | 66.67%/33.33% | 62.50%/37.50% | 0.091 | 0.763 |
| the only child (yes/no) | 58.33%/41.67% | 58.33%/41.67% | < 0.001 | 1.000 |
| political orientation (Trump/Hillary) | 66.67%/33.33% | 80.00%/20.00% | 0.978 | 0.323 |
|  | mean (SD) | Mean (SD) | Z value in  Mann-Whitney U test | *p* |
| educational level | 3.04 (0.46) | 3.17 (0.43) | -0.972 | 0.331 |
| household annual income per person (in 10,000 CNY) | 3.17 (1.27) | 3.12 (1.22) | 0.255 | 0.799 |
| birth sequence | 1.46 (0.93) | 1.46 (0.72) | -0.464 | 0.643 |
|  | mean (SD) | mean (SD) | t value | *p* |
| age | 21.75 (2.19) | 21.92 (3.76) | -0.187 | 0.852 |
| height (cm) | 174.00 (4.31) | 176.00 (5.60) | -1.387 | 0.172 |
| weight (kg) | 66.10 (8.57) | 67.94 (7.05) | -0.811 | 0.421 |
| number of siblings | 2.40 (1.07) | 2.00 (0.82) | 0.937 | 0.361 |
| duration of friendship (months) | 23.71 (13.86) | 22.96 (27.09) | 0.121 | 0.904 |
| frequency of contact (times per month) | 29.91 (25.11) | 29.53 (27.47) | 0.048 | 0.962 |
| friendship quality | 8.17 (0.95) | 8.04 (1.60) | 0.392 | 0.744 |
| significance of the friendship | 2.79 (0.41) | 2.63 (0.65) | 1.062 | 0.294 |
| considered by the friend in decision | 3.00 (0.66) | 2.85 (0.88) | 0.648 | 0.521 |
| considering the friend in decision | 2.88 (0.80) | 2.75 (0.79) | 0.521 | 0.605 |
| prior-experimental Positive and Negative Affect Schedule (Positive Affect) | 30.50 (5.95) | 32.54 (5.82) | -1.201 | 0.236 |
| prior-experimental Positive and Negative Affect Schedule (Negative Affect) | 19.50 (6.26) | 20.71 (8.07) | -0.580 | 0.565 |
| post-experimental Positive and Negative Affect Schedule (Positive Affect) | 30.08 (6.62) | 29.04 (7.21) | 0.521 | 0.605 |
| post-experimental Positive and Negative Affect Schedule (Negative Affect) | 20.04 (5.65) | 20.50 (8.18) | -0.226 | 0.822 |
| prior-experimental State Anxiety Inventory | 40.29 (9.56) | 38.79 (9.67) | 0.540 | 0.592 |
| post-experimental State Anxiety Inventory | 39.67 (9.49) | 41.08 (8.32) | -0.550 | 0.585 |
| Social dominance orientation | 54.30 (9.17) | 57.04 (17.44) | -0.667 | 0.508 |
| Big five neuroticism | 44.22 (10.13) | 42.33 (16.30) | 0.47 | 0.64 |
| Big five extraversion | 53.17 (9.49) | 50.04 (14.12) | 0.89 | 0.38 |
| Big five openness | 56.17 (7.73) | 54.00 (14.56) | 0.64 | 0.53 |
| Big five agreeableness | 55.83 (10.76) | 56.25 (13.88) | -0.12 | 0.91 |
| Big five conscientiousness | 55.57 (9.26) | 55.67 (15.79) | -0.03 | 0.98 |
| Interpersonal reactivity index perspective taking | 22.30 (2.01) | 22.61 (2.35) | -0.47 | 0.64 |
| Interpersonal reactivity index fantasy scale | 24.00 (3.57) | 23.91 (5.12) | 0.07 | 0.95 |
| Interpersonal reactivity index empathic concern | 24.52 (2.71) | 25.65 (3.23) | -1.29 | 0.21 |
| Interpersonal reactivity index personal distress | 21.30 (3.54) | 21.17 (3.99) | 0.12 | 0.91 |
| Toronto alexithymia total | 52.91 (8.78) | 51.48 (10.48) | 0.50 | 0.62 |
| Toronto alexithymia difficulty describing feelings | 14.52 (3.04) | 13.91 (3.36) | 0.64 | 0.52 |
| Toronto alexithymia difficulty identifying feeling | 19.61 (4.30) | 19.26 (4.95) | 0.25 | 0.80 |
| Toronto alexithymia externally oriented thinking | 19.78 (4.19) | 19.30 (4.35) | 0.38 | 0.71 |
| Barratt impulsiveness scale | 72.70 (6.42) | 68.17 (7.57) | 2.19 | 0.03 |
| Self esteem | 30.39 (4.15) | 28.91 (4.65) | 1.14 | 0.26 |
| Narcissistic personality inventory | 130.13 (30.60) | 127.00 (22.99) | 0.39 | 0.70 |
| Fear of negative evaluation | 95.48 (14.14) | 94.43 (18.01) | 0.22 | 0.83 |
| Individualism and collectivism horizontal individualism | 43.39 (5.63) | 40.96 (5.46) | 1.49 | 0.14 |
| Individualism and collectivism vertical individualism | 36.52 (4.39) | 35.30 (4.88) | 0.89 | 0.38 |
| Individualism and collectivism horizontal collectivism | 38.52 (7.35) | 38.87 (4.46) | -0.19 | 0.85 |
| Individualism and collectivism vertical collectivism | 36.48 (5.08) | 33.04 (5.38) | 2.23 | 0.03 |
| Adult attachment style closeness | 3.52 (0.85) | 3.61 (0.58) | -0.41 | 0.69 |
| Adult attachment style dependency | 3.22 (0.60) | 3.17 (0.49) | 0.27 | 0.79 |
| Adult attachment style anxiety | 2.91 (0.79) | 2.96 (0.88) | -0.18 | 0.86 |
| Adult attachment style closeness & dependency | 3.30 (0.64) | 3.22 (0.42) | 0.55 | 0.59 |
| Experiences in close relationships avoidance | 3.39 (0.99) | 3.22 (0.95) | 0.61 | 0.55 |
| Experiences in close relationships anxiety | 3.70 (1.15) | 3.74 (1.10) | -0.13 | 0.90 |
| Relationship scale secure | 3.04 (0.48) | 3.08 (0.65) | -0.24 | 0.81 |
| Relationship scale fearful | 2.91 (0.79) | 3.00 (0.93) | -0.34 | 0.73 |
| Relationship scale preoccupied | 2.96 (0.83) | 2.79 (0.78) | 0.71 | 0.49 |
| Relationship scale dismissing | 3.52 (0.67) | 3.54 (0.59) | -0.11 | 0.91 |
| Childhood trauma physical neglect | 7.22 (2.54) | 7.43 (3.22) | -0.25 | 0.80 |
| Childhood trauma emotional neglect | 9.87 (3.77) | 8.87 (3.55) | 0.93 | 0.36 |
| Childhood trauma sex abuse | 6.17 (2.19) | 5.74 (1.14) | 0.85 | 0.40 |
| Childhood trauma physical abuse | 7.09 (2.89) | 6.04 (1.85) | 1.46 | 0.15 |
| Childhood trauma emotional abuse | 8.26 (3.49) | 7.78 (2.49) | 0.54 | 0.60 |
| Father withdraw love | 25.00 (7.94) | 23.65 (8.17) | 0.57 | 0.57 |
| Mother withdraw love | 27.48 (10.25) | 25.48(9.96) | 0.67 | 0.51 |
| Intolerance of uncertainty scale | 77.35 (18.14) | 70.87 (18.60) | 1.20 | 0.24 |
| Autism quotient | 21.57 (5.65) | 22.38 (5.79) | -0.49 | 0.63 |
| Beck depression inventory | 3.58 (3.86) | 3.46 (4.05) | 0.11 | 0.91 |
| Buss-Perry aggression | 247.42 (54.05) | 238.88 (62.43) | 0.51 | 0.62 |
| Interaction anxiousness | 46.25 (9.43) | 45.29 (8.78) | 0.37 | 0.72 |
| Interpersonal trust | 69.58 (6.68) | 69.21 (9.13) | 0.16 | 0.87 |
| Trait anxiety inventory | 41.83 (7.45) | 42.67 (9.60) | -0.34 | 0.74 |
| Self-rating anxiety | 32.67 (6.16) | 34.21 (8.76) | -0.71 | 0.48 |
| Self-rating depression | 37.63 (7.00) | 39.42 (7.56) | -0.85 | 0.40 |
| UCLA loneliness | 42.67 (8.09) | 45.38 (8.19) | -1.15 | 0.26 |
| Happiness | 15.29 (6.33) | 14.63 (6.26) | 0.37 | 0.72 |
| Positive and negative reciprocity general | 41.88 (6.89) | 41.54 (7.00) | 0.17 | 0.87 |
| Positive and negative reciprocity Positive | 50.67 (5.83) | 49.75 (7.20) | 0.49 | 0.63 |
| Positive and negative reciprocity Negative | 37.38 (8.58) | 38.50 (8.33) | -0.46 | 0.65 |
| Social value orientation (degree) | 27.95 (15.53) | 28.08 (11.48) | -0.03 | 0.974 |
| Social value orientation (inequality aversion index) | 0.39 (0.23) | 0.43 (0.16) | -0.74 | 0.461 |
| Dictator game (gain) (amount of share) | 2.82 (1.59) | 3.21 (1.79) | -0.792 | 0.433 |
| Dictator game (loss) (amount of self loss) | 2.99 (1.76) | 3.26 (1.78) | -0.529 | 0.599 |
| Ultimatum game (gain) (amount of share) | 4.24 (0.78) | 4.47 (0.60) | -1.158 | 0.253 |
| Ultimatum game (loss) (amount of self loss) | 4.30 (0.87) | 4.43 (0.92) | -0.516 | 0.608 |
| Trust game with human (amount of invest) | 7.21 (2.21) | 6.96 (2.90) | 0.336 | 0.738 |
| Trust game with computer (amount of invest) | 6.71 (2.29) | 5.50 (3.28) | 1.478 | 0.146 |

PBO, placebo; AVP, arginine vasopressin; SD, standard deviation
